# Decoding mechanism of action and susceptibility to drug candidates from integrated transcriptome and chromatin state

**DOI:** 10.1101/2022.02.21.481294

**Authors:** Caterina Carraro, Lorenzo Bonaguro, Jonas Schulte-Schrepping, Arik Horne, Marie Oestreich, Stefanie Warnat-Herresthal, Tim Helbing, Michele De Franco, Kristian Händler, Sach Mukherjee, Thomas Ulas, Valentina Gandin, Richard Göttlich, Anna C. Aschenbrenner, Joachim L. Schultze, Barbara Gatto

## Abstract

Omics-based technologies are driving major advances in precision medicine but efforts are still required to consolidate their use in drug discovery. In this work, we exemplify the use of multi-omics to support the development of 3-chloropiperidines (3-CePs), a new class of candidate anticancer agents. Combined analyses of transcriptome and chromatin accessibility elucidated the mechanisms underlying sensitivity to test agents. Further, we implemented a new versatile strategy for the integration of RNA-seq and ATAC-seq data, able to accelerate and extend the standalone analyses of distinct omic layers. This platform guided the construction of a perturbation-informed basal signature able to predict cancer cell lines’ sensitivity and to further direct compound development against specific tumor types. Overall, this approach offered a scalable pipeline to support the early phases of drug discovery, understanding of mechanism and potentially inform the positioning of therapeutics in the clinic.

## Introduction

Omics technologies have revolutionized the classical *hypothesis-driven* paradigm of drug discovery, offering a new perspective for the systematic identification of targets and therapeutics.^1, 2^ An increasing number of examples are describing the use of these approaches to inspect the pharmacological profile of existing drugs, e.g. mechanism of action (MoA) and specific sensitivity biomarkers, as well as to assist their correct repositioning in the clinical practice.^3–6^ Compared to traditional approaches, omics-based methods capture the complexity of biological systems and pathological processes in its entirety at increasingly affordable costs.^3^ For this reason, refined strategies to handle the high-dimensional information of omics data are continuously investigated to expedite their routine use in drug development up to the clinics.^7, 8–14^

Recent works from our group highlighted 3-chloropiperidines (3-CePs) as a novel class of candidate anticancer agents developed to improve the pharmacological profile of nitrogen mustard-based chemotherapeutics.^15–21^ As intended, these agents were demonstrated to induce DNA lesions, a mechanism conceivably responsible for their cytotoxicity on tested cancer cell lines.^19–21^ Interestingly, despite their expected broad-acting MoA, a subset of derivatives showed a preferential activity against pancreatic adenocarcinoma BxPC-3 cells worth to be clinically translated, especially in light of the broad resistance of pancreatic tumors to most of the available treatments.^19–21^

The contribution of multi-omics to support early phases of drug discovery is growing exponentially in the era of precision medicine.^7^ Omics technologies have the potential to address some of the intrinsic difficulties of the traditional drug discovery and development path, assisting it early from target prioritization and hit identification up to the evaluation of candidates’ efficacy and safety.^4^ Drug-perturbation experiments have been employed to inspect the functionality of target proteins^22^ and the MoA of therapeutics, efficiently guiding the decision-making process in the development of lead compounds.^4^ The massive accumulation of genomic and transcriptomic profiles offers a precious substrate for the optimization of strategies able to predict susceptibility to known therapeutics^23–26^ refined by the continuous acquisition of data from high-throughput single-cell platforms.^10–12, 27, 28^ Beyond the widely used transcriptome analysis, changes in gene regulation can be evaluated in terms of chromatin accessibility by ATAC-seq.^29–31^ Examples of the joint use of these two omic techniques exist,^13, 14, 32, 33^ but their synergistic employment on compounds under early development is still underexplored.^3^

In this study, representative mono- (**M**) and bifunctional (**B**) 3-CePs bearing a single or double alkylating units (Fig. 1 A) were selected to exemplify the use of a multi-omic approach to investigate the molecular determinants of susceptibility to novel drug candidates and their MoA.^19–21^ We analyzed transcriptional changes and chromatin status upon treatment in a high- (pancreatic adenocarcinoma BxPC-3) and low-sensitive (colorectal adenocarcinoma HCT-15) cancer cell lines by RNA-seq and ATAC-seq.^29–31^ In addition, we implemented our multi-omics pipeline in drug discovery to derive perturbation-informed signatures predicting compound sensitivity. Overall, the proposed approach not only allowed to identify potentially more susceptible target tumor types for the further development of test compounds, but also offered a versatile predictive framework to support precision oncology in a clinical setting.

**Figure 1.**
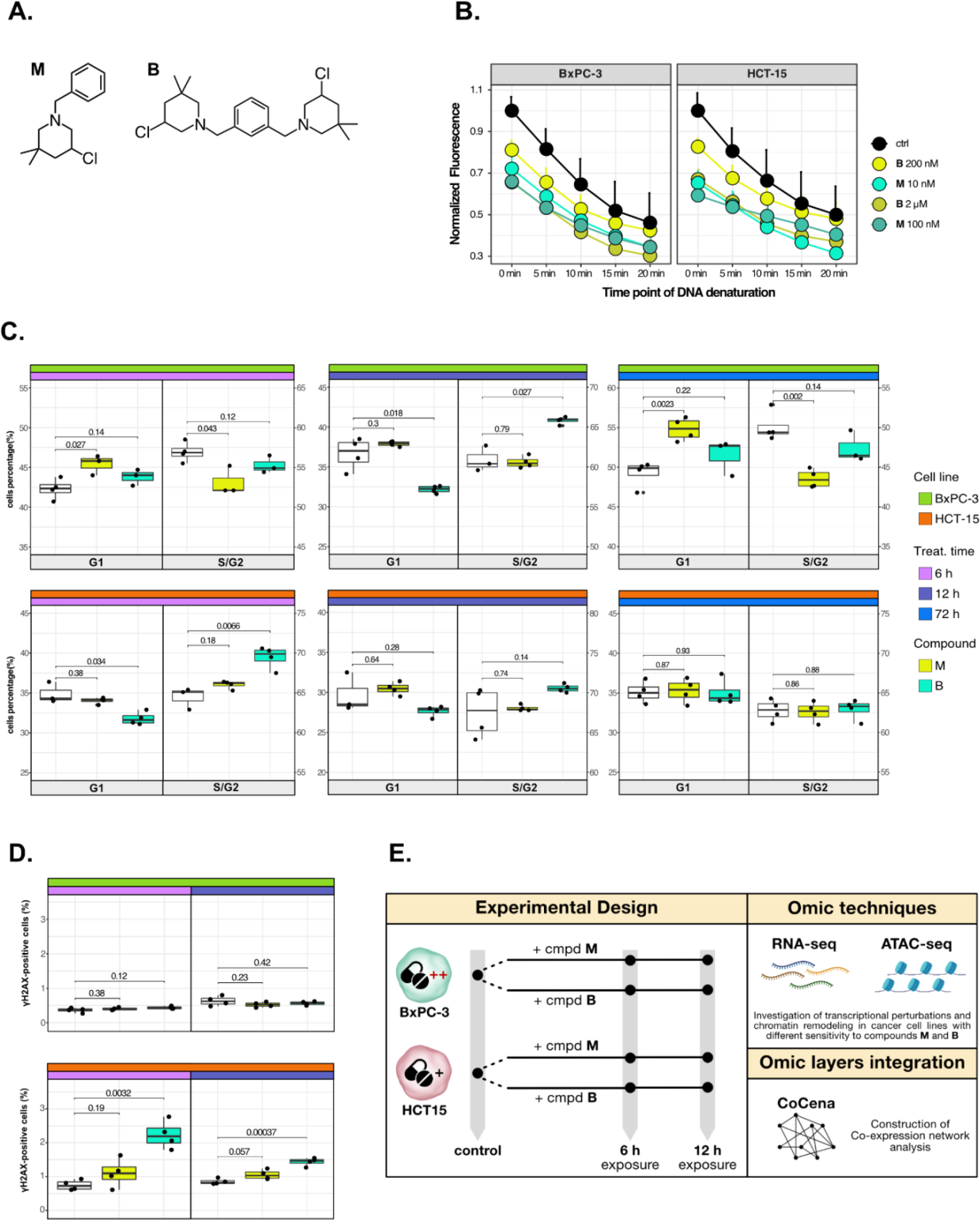
Cancer tropism of 3-CePs is not explained by DNA damage. **A** Chemical structure of the analyzed 3-CePs (M = monofunctional, B = bifunctional). **B** Quantification of genomic DNA damage in BxPC-3 and HCT-15 cells treated with M (10 nM and 100 nM), B (200 nM and 2 µM) or DMSO 0.5% (ctrl) for 6 h and analyzed by the Fast Micromethod single-strand-break assay: alkaline denaturation of DNA is followed in time up to 20 min by monitoring the fluorescence of the dsDNA-specific PicoGreen dye. **C** Cell cycle distribution (accumulation in G1 vs G2/S phases) of BxPC-3 and HCT-15 cells treated with M (10 nM), B (200 nM) or DMSO 0.5% for 6 h, 12 h and 72 h analyzed by FACS. Three biological replicates were obtained per condition and unpaired two-tailed Student’s *t*-test was performed to assess statistical significance (p < 0.05). **D** Analysis of H2AX phosphorylation in BxPC-3 and HCT-15 cells treated with M (10 nM), B (200 nM) or DMSO 0.5% for 6 h and 12 h analyzed by FACS. Three biological replicates were obtained per condition and unpaired two-tailed Student’s *t*-test was performed to assess statistical significance (p < 0.05). **E** Schematic representation of the adopted omic-based approach.

## Results

### Cancer tropism of 3-CePs is not explained by DNA damage

The mono- (**M**) and bifunctional (**B**) 3-chloropiperidines (3-CePs, Fig. 1 A), despite having different potencies, were shown to be particularly active against BxPC-3 pancreatic adenocarcinoma cells.^19, 20^ From this premise, the two compounds were selected along with the highly sensitive BxPC-3 cell line and the low-sensitive HCT-15 colorectal adenocarcinoma one to illustrate how integrative omics approaches unveil the molecular mechanisms responsible for the described cellular tropism.

First, to assess whether 3-CePs-induced DNA damage itself would differ in the two cell lines upon treatment, we measured the accumulation of DNA single-strand breaks after 6 h of treatment with both compounds at their cytotoxicity IC50s in BxPC-3 and at a ten-times higher concentration (10 nM and 100 nM **M;** 200 nM and 2 µM **B**).^34^ Surprisingly, the two cell lines showed very comparable DNA damage accumulation, in both cases higher after treatment with **M** compared to **B** (Fig. 1 B). These results clearly pointed towards differential responses in the two cell lines downstream of DNA damage.

Since alkylating agents are known to alter the progression of the cell cycle,^35–37^ we next performed a cell cycle distribution analysis by flow cytometry after different times of treatment (6 h, 12 h, 72 h) with both compounds (Fig. 1 C). While **M** induced a persisting block in G1 throughout the observation time in BxPC-3 cells, this block was absent in HCT-15 cells. In contrast, **B** induced an early G2/S block in HCT-15 cells (6 h), which was not observed at later time points, while such a block was most obvious at 12 h for BxPC-3 cells. Despite similar DNA damage accumulation, these findings clearly indicated a different behavior for the two cancer cell lines in terms of cell cycle progression after treatment with the two 3-CePs.

To determine additional mechanisms explaining differential sensitivity to 3-CePs, we measured the activation of the DNA repair machinery as another key aspect in the cellular response to genotoxicants.^38^ To verify the ability of the two cancer cell lines to detect double-strand breaks (DSBs), we assessed the phosphorylation of H2AX (γH2AX), an early event of the DNA damage response (DDR),^39^ by flow cytometry after 6 h and 12 h of treatment with both agents (Fig. 1 D). Interestingly, despite the comparable DNA damage accumulation in the two cell lines, only HCT-15 showed an increase in the γH2AX-positive population, suggesting a more efficient engagement of the DNA repair machinery.

Taken together, these results indicated that cell-specific mechanisms after the first event of DNA damage are responsible for the different susceptibilities to 3-CePs.

### Treatments elicit cell-specific transcriptional changes

Different genetic and epigenetic factors define the responsiveness of tumor cells to chemotherapeutic agents.^40^ To address these globally, we analyzed changes in the transcriptome of the high- and low-sensitive cell lines after treatment with the two 3-CePs (Fig. 1 E). RNA-seq was performed on total RNA of HCT-15 and BxPC-3 cells exposed to DMSO 0.5% (control) or treated with **M** (10 nM) or **B** (200 nM) for 6 h and 12 h (Fig. 2 A, S2 A) as in previous experiments.

**Figure 2.**
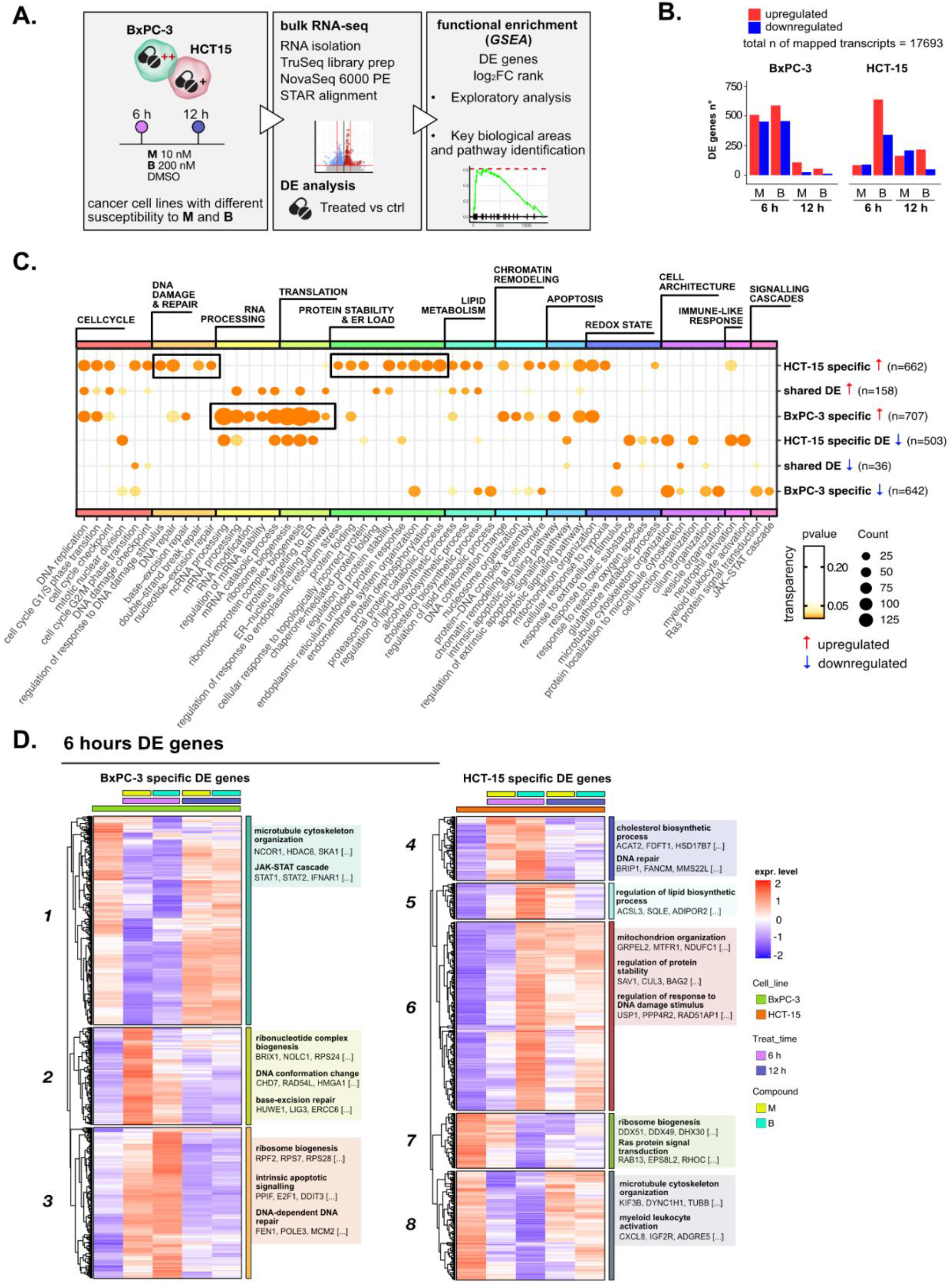
Treatments elicit cell-specific transcriptional changes. **A** Overview of the applied workflow for the RNA-seq analysis. **B** Number of up- (red) and downregulated (blue) DE genes in BxPC-3 and HCT-15 cells after treatment with M (10 nM), B (200 nM) or DMSO 0.5% (ctrl) for 6 h and 12 h (adjusted p threshold = 0.05, shrinkage = TRUE, multiple testing method = IHW). **C** GO database functional enrichment (GSEA) on cell-specific and shared up- and downregulated DE genes. For each identified biological process, enrichments in terms of Count and p-value of representative terms are reported (p < 0.05). **D** Expression level of cell-specific 6 h DE genes across test conditions. GSEA was performed on modules with similar regulation identified by hierarchical clustering: for each cluster, representative GO terms and genes of the associated load are reported.

Principal component analysis (PCA) of all transcripts separated samples within each cell line according to treatment and time-point (Fig. S2 B), suggesting a clear transcriptional reprogramming after treatment. In fact, differential expression (DE) analysis pointed out that the expression of a large number of genes changed significantly in both cell lines after exposure to 3-CePs (Fig. 2 B, S2 C), especially at 6 h in BxPC-3 cells and upon treatment with **B** in HCT-15 cells.

Gene Ontology (GO) enrichment was performed on the DE genes to determine signaling pathways and transcriptional programs explaining the observed differences. In a first explorative approach, we generated the union of DE genes per cell line irrespective of compound and time point, which allowed us also to distinguish between cell type-specific or shared DE genes (Fig. S2 D). The most representative biological processes identified by this analysis (Fig. S2 E, Supplementary data 1) are reported in Fig. 2 C (see *Methods* and Fig. S2 F for further details).

Unexpectedly, we identified a strong translational response in BxPC-3 cells after treatment, a process which is typically attenuated in stress conditions, as was the exposure to our DNA damaging agents, to allow proper recovery of the protein quality control machinery.^41, 42^ In contrast, a strong regulation of genes mediating protein stability and catabolism was observed in the low-sensitive cell line. In addition, HCT-15 cells activated genes involved in the DDR, consistently with their higher ability to detect and respond to DSBs. Both these two mechanisms pointed towards the activation of an adaptive stress response in the low-sensitive cell line.

To further characterize these transcriptional changes over time in a cell type-specific context, we grouped the DE genes at 6 and 12 h in modules according to the similarity in their expression profiles and performed a functional enrichment on genes with similar expression patterns (Fig 2 D and S3 A, Supplementary data 2). Genes involved in ribosome biogenesis and DNA repair turned out to be upregulated particularly after 6 h of treatment in BxPC-3 cells (Clusters *2* and *3*, Fig. 2 D). Besides, silencing of pro-survival genes involved in microtubule organization and the JAK-STAT cascade (Cluster *1*, Fig 2 D) was detected at the same time point. Only after 12 h of treatment (Fig. S3 A), BxPC-3 cells boosted carbohydrate metabolism, most likely an attempt to recover *in extremis*.^43^

Also HCT-15 cells upregulated clusters of genes mediating DNA repair, protein stability and mitochondrial activity as early as 6 h of treatment, suggesting this time point as the most informative to describe the response to 3-CePs (Clusters *4* and *6*, Figure 2 D). In contrast to BxPC-3 cells, HCT-15 downregulated genes involved in translation and ribosome biogenesis from 6 h of exposure (Cluster *7*, Figure 2 D), while intensifying their response to oxidative stress after 12 h (Cluster *17*, Figure S3 A).

This exploratory analysis showed clearly different transcriptional responses and distinct time dynamics in BxPC-3 compared to HCT-15 cells, most likely responsible for their different susceptibility to 3-CePs. In particular, our findings pointed towards DNA repair and proteostasis as key mechanisms tuning sensitivity to the compounds, as further confirmed by inspecting the complete rank of DE genes *via* gene set enrichment analysis (GSEA, Fig. S3 B).^44^

### DNA repair and proteostasis are key modulators of the response to 3-CePs

For their key role in the response to 3-CePs, DNA repair and protein homeostasis were further analyzed to clarify their contribution to BxPC-3 sensitivity.

Interestingly, DNA repair was activated in both cell lines early after 6 h of treatment but with a different modulation (Fig. 3 A). First, base-excision repair (BER) was suggested as the preferential pathway of BxPC-3 by GO enrichment while HCT-15 relied mostly on nucleotide-excision repair (NER), unleashing a generally stronger activation of the DDR. In detail, HCT-15 DE genes contributing to the response to the DNA damage stimulus were strongly upregulated already after 6 h especially in response to **B**, while activated only after 12 h in BxPC-3 (Fig. 3 B). In contrast, genes such as *PPP4R2* and *RAD51AP1*, both involved in the first phases of DSBs repair,^45, 46^ were even downregulated in BxPC-3 cells at 6h.

**Figure 3.**
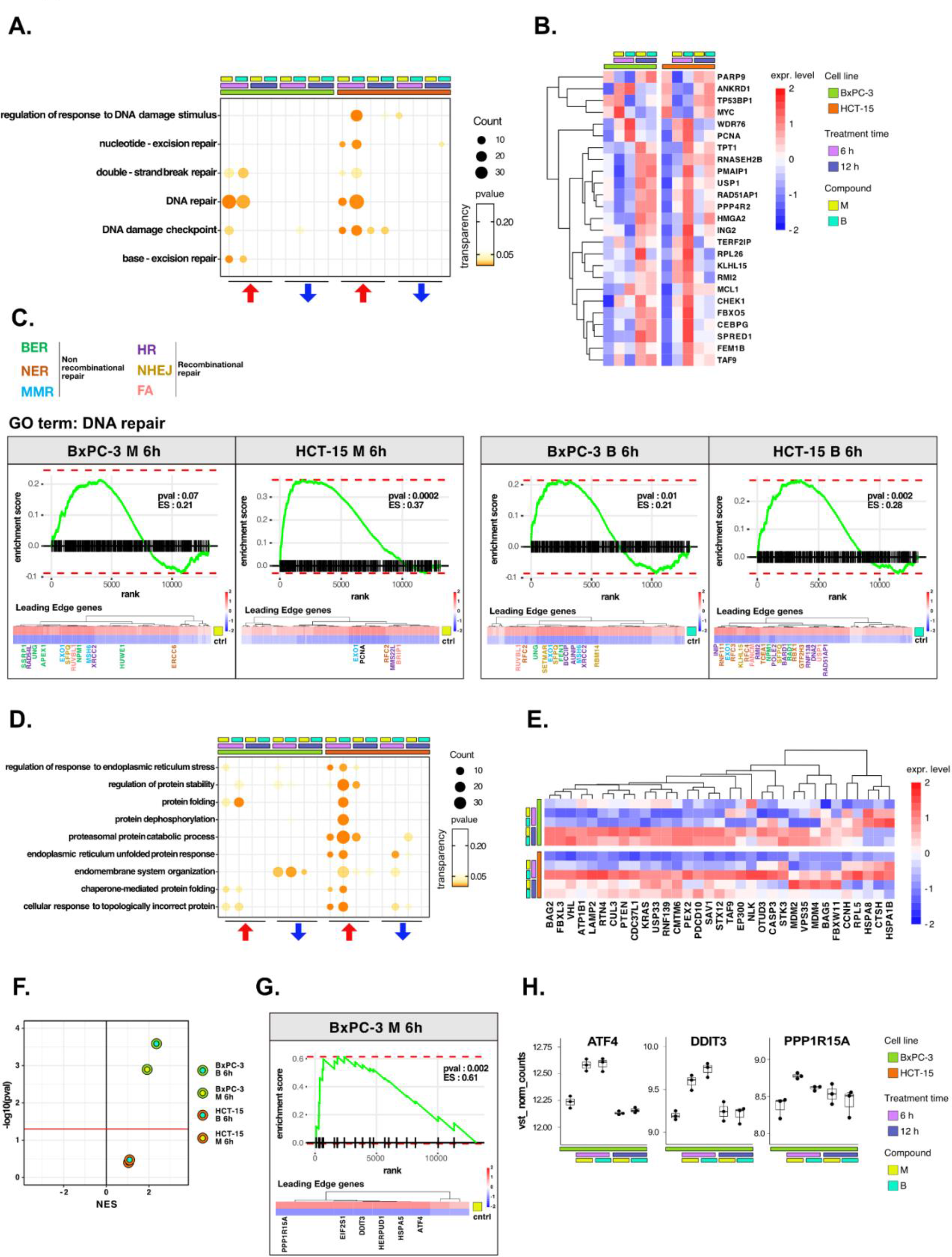
DNA repair and proteostasis are key modulators of the response to 3- CePs. **A** GSEA for terms related to *DNA damage and repair* performed on DE genes detected in each of the considered treated vs control comparisons. For each GO term (p < 0.05), enrichments in terms of Count and p-value are reported. **B** Expression level of DE genes included in the load of the GO term *regulation of response to DNA damage stimulus* (HCT-15 cells, p < 0.05) in BxPC-3 and HCT-15 cells. **C** GSEA enrichment plots for the *DNA repair* pathway obtained from log2FC ranks for each of the considered treated vs control comparisons. The expression of leading edge genes is also shown, where key DE genes are reported with the same color of their associated DNA repair pathways (BER=base excision repair, NER=nucleotide-excision repair, MMR=mismatch repair, HR=homologous recombination, NHEJ=non-homologous end joining, FA=Fanconi anemia pathway).^47–54, 82, 95, 96, 114–133^ **D** GSEA for terms related to *protein stability and ER load* performed on DE genes detected in each comparison. For each GO term, enrichments in terms of Count and p-value are reported. **E** Expression level of DE genes included in the load of the GO term *proteasomal protein catabolic process* (HCT-15 cells, p < 0.05) in BxPC-3 and HCT-15 cells. **F** NES (normalized enrichment score) and - log10pval for the log2FC rank-based GSEA enrichment of the GO term *PERK-mediated UPR* in treated vs control comparisons. **G** GSEA enrichment plot for the *PERK-mediated UPR* pathway obtained from log2FC rank in the M 6 h vs control comparison in BxPC-3 cells. The expression of leading edge genes is also shown, where key DE genes of the mentioned pathway are reported. **H** Boxplots showing the expression level of ATF4, DDIT3 and PPP1R15A (vst-transformed normalized counts) in BxPC-3 cells.

The more efficient activation of DNA repair in HCT-15 was further confirmed on the overall rank of genes by GSEA at 6 h of treatment (Fig. 3 C). As anticipated, most of the DE genes leading the enrichment in HCT- 15 belonged to NER (e.g. *GTF2H3*, *RBX1*) and other recombinational pathways such as Homologous Repair (HR) (e.g. *MMS22L*, *BARD1*) and Fanconi Anemia (FA) (e.g. *BRIP1*, *FANCM*), all better suited for the efficient repair of bulky lesions and highly toxic DSBs and crosslinks.^47–52^ On the other hand, DE genes in BxPC-3 cells were mostly related to BER (e.g. *APEX1*, *UNG*) and MMR (Mismatch Repair) (e.g. *MSH6*, *EXO1*), which contribute to the repair of smaller lesions and mismatches.^53, 54^

In the analysis, proteostasis was identified as a second key biological process strictly related to genotoxic stress.^55, 56^ HCT-15 cells engaged the protein folding and catabolism apparatus in response to 3-CePs, especially to **B** already at the early time point (Fig. 3 D). As observed for DNA repair, DE genes contributing to protein catabolism were upregulated as early as 6 h of exposure in HCT-15 cells, while even downregulated at the same time point in BxPC-3 and only upregulated after 12 h (Fig. 3 E). This response involved chaperones and co-chaperones (e.g. *HSPA8*, *HSPA1B*, *BAG2*, *BAG5*), other genes mediating protein catabolism (e.g. *LAMP2*, *CUL3*) and ER morphogenesis (e.g. *RTN4*).^57–60^ Interestingly, a transcriptional pattern revealed by GSEA at 6 h of treatment highlighted an intense positive modulation of the PERK-mediated branch of the unfolded protein response (UPR) specifically in BxPC-3 (Fig. 3 F). Even more enlightening were the DE genes leading the enrichment: *ATF4*, *DDIT3* (CHOP) and *PPP1R15A* (GADD34) were significantly upregulated after 6 h of exposure only in this cell line (Fig. 3 G, H). These genes participate in the PERK-mediated UPR triggering cell death after prolonged ER stress through the aberrant recovery of translation, which induces proteotoxicity.^61, 62^ This mechanism would reasonably explain the ribosome biogenesis signature observed in BxPC-3 cells. Consistently, recent work reported a particular susceptibility for pancreatic cancer adenocarcinoma to ER stress and protein dyshomeostasis.^63^

Furthermore, the ability of HCT-15 cells to control proteostasis may also depend on the activation of lipid and cholesterol biosynthesis in response to the compounds (Fig. S4 A). In fact, among other known pro-survival functions, these pathways contribute to resolving ER stress through pathways involving e.g. the Stearoyl-CoA Desaturase (*SCD*) enzyme, for which we detected a significant upregulation of the respective transcript in HCT-15 (Fig. S4 B).^64, 65^

Overall, the transcriptome analysis of this *in vitro* perturbation experiment allowed us to dissect the different responses to 3-CePs in our model cell lines, pointing towards protein homeostasis and DDR imbalances as mechanisms responsible for the high susceptibility of BxPC-3 cells.

### The response to 3-CePs is further regulated at the chromatin level

The transcriptome analysis unveiled a defined framework of responses tuning the sensitivity to 3-CePs. To further characterize them at the epigenetic level, we examined chromatin accessibility in nuclei of BxPC-3 and HCT-15 cells treated with **M** and **B** for 6 h and 12 h (Fig. 4 A, Fig. S5 A) by ATAC-seq.

**Figure 4.**
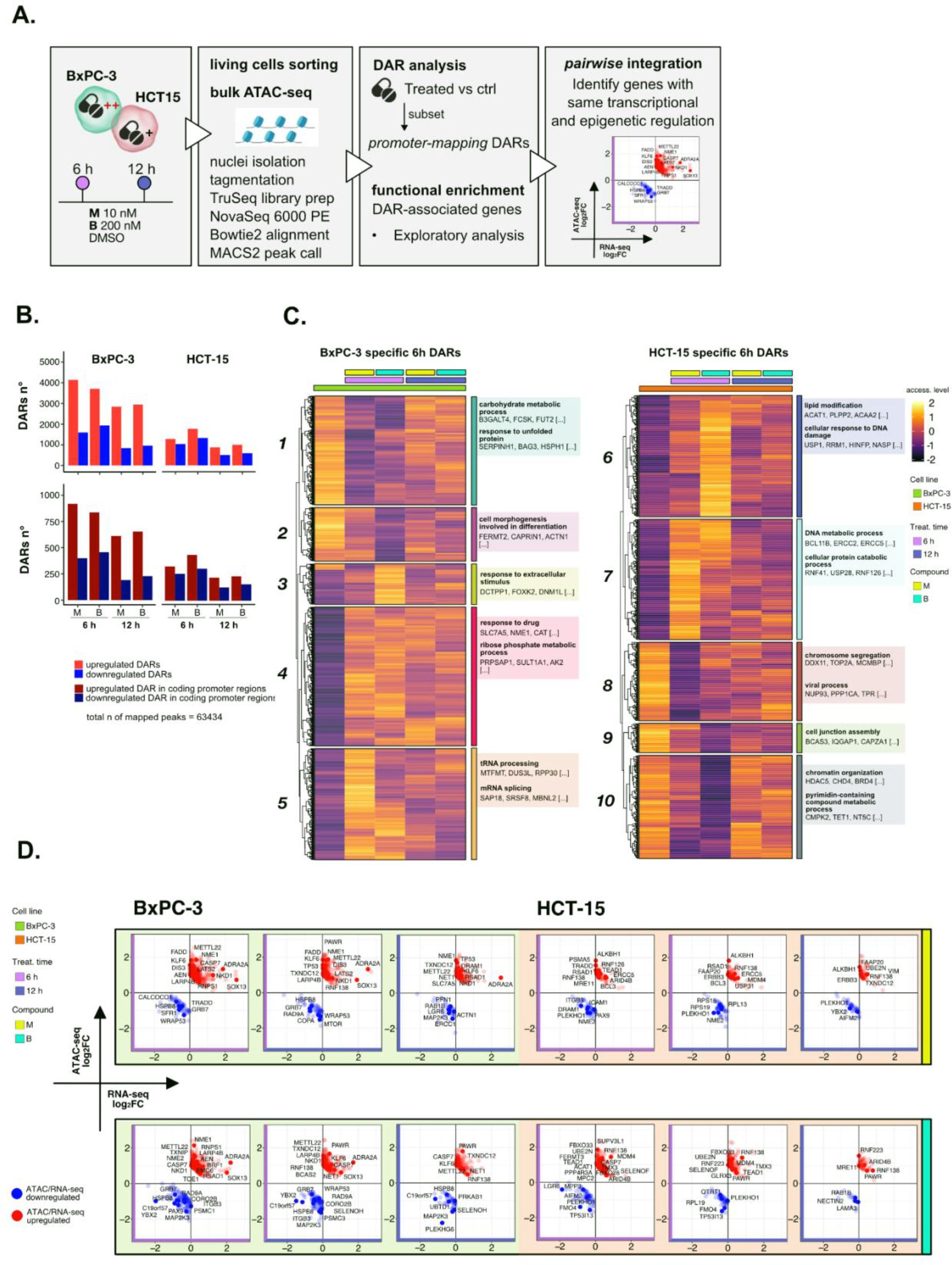
The response to 3-CePs is further regulated at the chromatin level. **A** Overview of the applied workflow for the ATAC-seq analysis. **B** Number of up- (red) and downregulated (blue) DARs in BxPC-3 and HCT-15 cells after treatment with M (10 nM), B (200 nM) or DMSO 0.5% (ctrl) for 6 h and 12 h (p-value threshold = 0.05, shrinkage = TRUE). Light blue/red = all detected DARs, dark blue/red = protein coding DARs mapping in promoter regions. **C** Accessibility level of cell-specific 6 h DARs across test conditions. GSEA was performed on genes associated with DARs with similar regulation, grouped in modules identified by hierarchical clustering: for each cluster, representative GO terms and genes of the associated load are reported. **D** Pairwise integration: ratio-ratio plots report the RNA-seq and ATAC-seq log2FCs of genes showing the same direction of transcriptional and chromatin accessibility regulation. Integration was performed not only at the same time point in both omic layers, but also between chromatin changes at 6 h and transcriptional responses at 12 h.

3-CePs induced evident epigenetic changes in both cell lines, as suggested by PCA (Fig. S5 B) and confirmed by the number of differentially accessible regions (DARs) identified especially in BxPC-3 cells (Fig. 4 B, Fig. S5 C). For further downstream analyses we focused on DARs mapping to promoters, whose specific condensation or compaction contribute to modulation of transcription of associated genes (Fig. 4 B).

Also in this case, to better describe the timing of chromatin remodeling, cell-specific promoter-associated DARs elicited after 6 h and 12 h of treatment were grouped in clusters sharing a similar pattern of regulation and functional enrichment was performed on the associated genes (Fig. 4 C, S5 D, Supplementary data 3).

In BxPC-3 cells, we observed condensation of promoters involved in carbohydrate metabolism and others mediating protein folding and UPR after 6 h of exposure (Cluster *1*, Fig. 4 C), most likely contributing to the transcriptional downregulation of such processes observed at the same time point.^57, 66^ On the contrary, relaxation of peaks involved in tRNA metabolism and mRNA splicing were detected, in line with the upregulation of translation and RNA processing evidenced by RNA-seq. In HCT-15 cells, relaxation of promoters involved in the DDR, lipid metabolism (Cluster *6*, Fig. 4 C) as well as protein catabolism (Cluster *7*, Fig. 4 C) was observed, again in line with our observations on transcriptome level. Altogether, these results attested that the regulation of elicited transcriptional pathways was accommodated by changes at the chromatin level, adding new information on the possible mechanisms determining the cellular responses to 3-CePs.

A critical step in the analysis of multi-omic datasets is the integration of information obtained from the different layers. Though valuable strategies have been developed in recent years to integrate RNA-seq and ATAC-seq data, alternatives are still required to optimize and enlarge the functional information obtained from the combination of these powerful techniques.^67–69^ In this study, we approached data integration through two alternative strategies, that we called *pairwise* and *crosswise*.

As a first level of integration, we identified genes with concordant regulation in RNA-seq and ATAC-seq upon treatment. In this *pairwise* integration, we compared the direction of transcriptional regulation of genes to the accessibility of their promoters, as specified in the *Methods* section and shown in Fig. 4 D. Given the biological delay that could exist between chromatin remodeling and a detectable variation in transcript level, pairwise comparisons were also considered between chromatin changes after 6 h and transcriptional responses after 12 h of treatment.

Among genes with coherent regulation in BxPC-3 cells we found the tumor suppressors *ADRA2A*, *NME1,* and *KLF6* to be upregulated, elicited after treatment with both agents, and *LATS2* and *NME2* specific for **M** and **B**, respectively.^70–73^ Besides, other genes were involved in translation and RNA processing such as *RNPS1* and *LARP4B*,^74, 75^ and apoptosis such as *AEN*, *PAWR,* and *CASP7*.^76–78^ Interestingly, BxPC-3 also negatively regulated *TXNIP*, an inhibitor of the oxidative stress regulator thioredoxin, after treatment with **B**.^79^ Conversely, among downregulated hits we found apoptosis inhibitors such as *WRAP53* and *TRADD*, as well as *HSPB8*, *CALCOCO1* and *SELENOH*, all involved in the resolution of ER and oxidative stress.^78, 80, 81^ In HCT-15 cells, among identified positively regulated genes some were involved in DNA repair such as *MRE11*, *MDM4*, *RNF138*,^82, 83^ others were oncogenes such as *VIM* and *ARID4B* or apoptosis inhibitors like *TRADD*.^84, 85^ Notably, some genes involved in the modulation of the redox balance (*GLRX3*, *SELENOF*) showed double regulation after treatment with **B** as well as others active in proteostasis (*PSMA5* after exposure to **M**, *UBE2N* to **B**).^81, 86^ Among the downregulated genes, some were associated to cell adhesion (*PLEKHO1*, *ITGB3*, *ICAM1*) and translation (*RPL19*, *RPL13*).^87^

Collectively, *pairwise* integration of RNA-seq and ATAC-seq shed light on genes with robust regulation at the transcriptional and chromatin level, adding further details to the previously identified response pathways.

### Crosswise integration expedites the comprehension of multi-omic data

Through the *pairwise* approach, we identified genes with both transcriptional and chromatin regulation which significantly contributed to the observed cellular response. We further evaluated the crosstalk between RNA-seq and ATAC-seq at a different level by focusing on groups of genes co-regulated in the two omic layers. The identification of genes sharing similar regulation across conditions either at the transcriptional or chromatin level would maximize the detection of interacting pathways and regulatory processes, e.g. as a result of chromatin changes in promoters tuning the transcription of a certain gene set. This approach, which we termed *crosswise* integration, was achieved by vertical Construction of Co-expression network analysis (vCoCena). vCoCena is designed to define modules of genes and/or genomic markers such as DARs with a similar pattern of regulation across conditions in multiple omic datasets. As a first step, we created separate co-expression networks for the RNA-seq and ATAC-seq layers (Fig. 5 A, S6 A). To prevent the construction of a network mostly describing the difference between the two cell lines, we first calculated separate networks for BxPC- 3 and HCT-15 cells which were then integrated horizontally (hCoCena).^88^ The union of all DE and promoter DAR-associated genes detected in treated conditions was selected as input for constructing all networks. Clustering of the resulting RNA-seq and ATAC-seq networks identified a relevant number of gene modules with highly specific regulatory patterns (Fig. S6 B and C). At this point, the vertical, inter-omic integration (vCoCena) was applied to construct the final network consolidating the information from transcriptome and chromatin accessibility (Fig. S6 D, see *Methods* for details). The new network was then reclustered resulting in integrated modules of co-regulation including nodes originally derived from the two separate layers in different ratios, as shown in Fig. 5 B.

**Figure 5.**
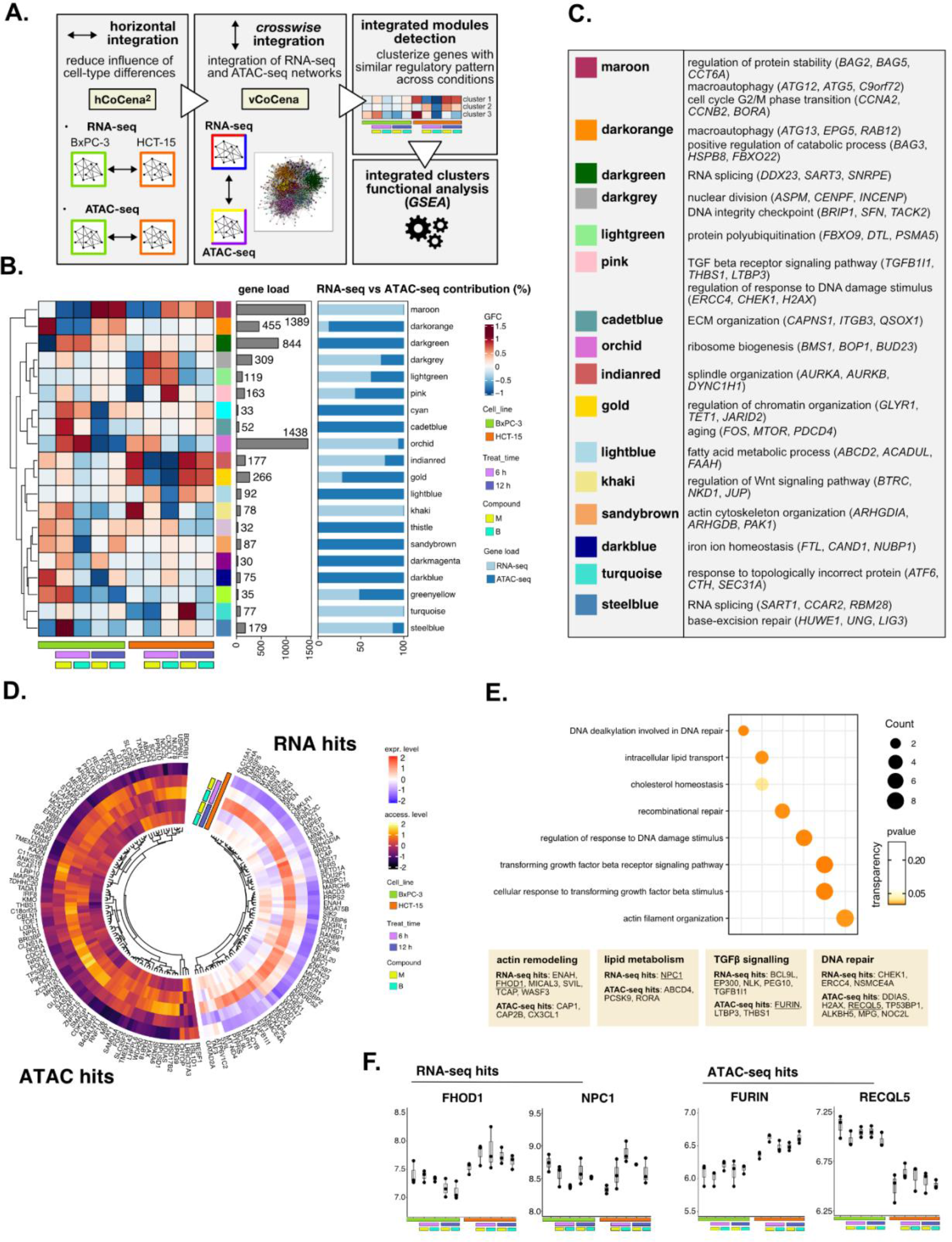
Crosswise integration expedites the comprehension of multi-omic data. **A** Overview of the applied workflow for the *crosswise* integration analysis. **B** Integrated modules of genes from the RNA-seq and ATAC-seq layers obtained with vCoCena and associated GFC (group fold change) pattern of regulation across conditions. The relative contribution of hits from the RNA-seq or ATAC-seq layers is also reported for each module. **C** Representative GO terms (p < 0.05) for the most relevant modules of genes, identified by GSEA. Enrichments in terms of Count and p-value are reported. **D** Expression and chromatin accessibility levels in HCT-15 cells of genes included in the *pink* module (nodes can come from the RNA-seq or ATAC-seq layer). **E** Most representative GO terms from GSEA on genes of the *pink* module (key areas: actin remodeling, lipid metabolism, TGF β signaling, DNA repair). For each GO term (p < 0.05), enrichments in terms of Count and p-value are reported. **F** Boxplots showing the expression level of FHOD1, NPC1, FURIN and RECQL5 (vst-transformed normalized counts) in BxPC-3 and HCT-15 cells.

The approach combined genes sharing similar regulation in the respective omic dataset, as approximated by the GFC pattern, with the postulate that genes grouped together cooperate in specific cellular processes. To define the underlying mechanisms, GO enrichment was performed on genes included in each of the modules and representative biological terms for the most relevant clusters were reported in Fig. 5 C (Supplementary data 4). Some modules validated the information obtained through previous analyses (Fig. S6 E): both the *maroon* and *darkorange* clusters suggested macroautophagy as a putative pathway accounting for the enhanced catabolism observed in HCT-15 cells.^89^ Consistently, the former RNA-seq-based module was downregulated at 6 h in BxPC-3 but upregulated already after 6 h with **B** in HCT-15, while the latter ATAC-seq-based module included peaks condensing after 6 h only in BxPC- 3,confirming the latter cell line as refractory to a rapid engagement of its protein catabolism apparatus. Another mostly RNA-seq-based module validating our previous approach was the *orchid* module, upregulated after 6 h in BxPC-3, containing genes involved in ribosome biogenesis. The *darkgrey* cluster instead, more balanced in terms of contribution from the two omic layers, showed positive regulation only in HCT-15 cells and included hits involved in DDR.

However, the *crosswise* integration also identified additional regulation, exemplified by the *pink* module. As approximated by the associated GFCs pattern, its 163 genes were positively modulated only in HCT-15 cells especially after 6 h of treatment with **B** (Fig. 5 D). Interestingly, functional enrichment identified hits both from RNA-seq and ATAC-seq involved in actin remodeling (Fig. 5 E), a mechanism affecting morphology and function of cancer cells (e.g. *FHOD1*, Fig. 5 F).^90–92^ Other module genes, such as *FURIN*, positively regulated at the chromatin level (Fig. 5 F), belonged to TGFβ signaling (Fig. 5 E), an emerging player in cancer drug resistance.^93, 94^ In addition, the module included genes of lipid metabolism and DNA repair belonging to both omic layers, which was in line with our initial findings (Fig. 5 E and F).

Overall, the *crosswise* integration of RNA-seq and ATAC-seq data allowed an efficient combination of the functional information from the two omics layers. Clearly, this approach added further biology to what we had identified when analyzing transcriptional and chromatin landscape regulation individually.

### Perturbation-informed basal signatures efficiently predict sensitivity to our candidate drugs

The information derived from the *crosswise* integration was employed to construct a signature of sensitivity to 3-CePs. Being more potent, **M** was selected as reference to describe a sensitivity prediction framework based on the use of a perturbation-informed omic signature (Fig. 6 A, S7 A, Methods).

**Figure 6.**
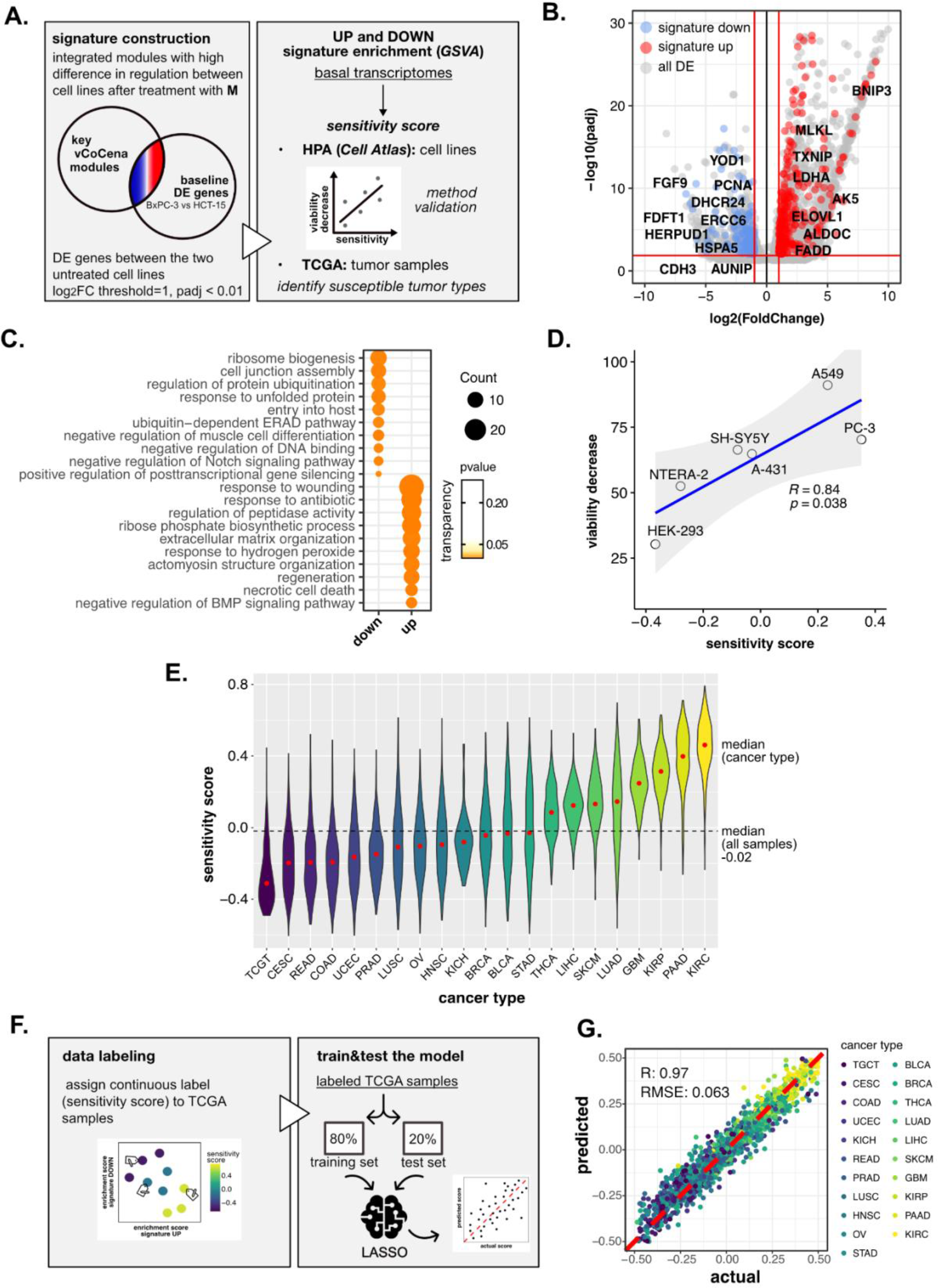
Perturbation-informed basal signatures efficiently predict sensitivity to our candidate drugs. **A** Overview of the applied workflow for the sensitivity signature construction and associated drug susceptibility prediction. **B** M sensitivity signature genes (red = signature up, blue = signature down) pinpointed from all DE genes in the BxPC-3 vs HCT-15 baseline comparison. **C** Representative GO terms (p < 0.05) for genes of the M sensitivity signature (up and down), identified by GSEA. Enrichments in terms of Count and p-value are reported. **D** Pearson correlation between predicted sensitivity score and viability decrease in a subset of HPA (*Cell Atlas*) cell lines (validation set). **E** Sensitivity scores predicted from GSVA enrichment of our up and down signatures in RNA-seq profiles of TCGA tumor samples. Median values for all sample scores and within each tumor type are reported. **F** Overview of the applied workflow for the LASSO-based ML setup. **G** Predictive outcome of the trained model (Pearson correlation R and RMSE are reported).

First, we selected vCoCena clusters with a marked difference in regulation between the two cell lines after treatment with **M,** considering only the most informative time point of 6 h (selected modules: *cyan*, *darkgreen*, *darkgrey*, *darkorange*, *gold*, *indianred*, *khaki*, *lightgreen*, *steelblue*, *orchid*; module selection criteria are described in detail in the *Methods* section). According to our analysis, genes that belong to these modules, coming both from RNA-seq and ATAC-seq analyses, are expected to be the major determinants of the differential susceptibility in the two cell lines.

Importantly, we postulated that features accounting for sensitivity should be intrinsic for the two cell lines, thus explained already by significant differences in their basal status. For this reason, we performed DE analysis between untreated BxPC-3 and HCT-15 control groups, identifying genes up- and downregulated at the transcriptional level in the high-sensitive cell line, and sorted out only those belonging to previously selected modules. This approach resulted in a subgroup of genes with different basal expression in BxPC-3 cells as well as a sufficiently compound- and cell line-specific regulation upon perturbation. This perturbation-informed signature was composed of 294 genes upregulated (signature *up*) and 170 genes downregulated (signature *down*) in the high-sensitive BxPC-3 cells (Fig. 6 B, gene list available in Supplementary data 5). GO enrichment on these genes identified protein synthesis, folding and catabolism, as well as cell adhesion, matrix organization and actin remodeling among the most significant biological functions (Fig. 6 C). Some interesting genes in the *up* signature were *BNIP3* and *FADD*, both proapoptotic, as well as *TXNIP*, already identified as a thioredoxin inhibitor. Among those composing the *down* signature, we identified *YOD1*, *HERPUD1* and *HSPA5*, involved in protein homeostasis and ER stress, but also *ERCC6* and *AUNIP* of the DDR (Fig. 6 B).^95–98^

To determine the robustness of the obtained signature and its ability to predict sensitivity to **M**, we next performed a gene set variation analysis (GSVA) on publicly available transcriptomes of common cell lines,^99^ testing for both the *up* and *down* signatures (Fig. S7 B). A sensitivity score was calculated for each cell line as the difference between the enrichment scores (ES) of the *up* and the *down* signatures. The predicted rank was validated experimentally on representative cell lines (A-431, A549, HEK- 293, NTERA-2, PC-3, SH-SY5Y) demonstrating the strong predictive capacity of our perturbation-informed signature (Pearson’s R=0.84, p=0.038, Fig. 6 D). This signature outperformed a random one containing the same number of genes (R=-0.48, p=0.34, Fig. S7 C and D) and also a signature of equal size composed by the top up- and downregulated genes between the two cell lines (R=0.34, p=0.51, Fig. S7 E and F, gene list available in Supplementary data 6). Collectively, our crosswise integration approach resulted in a perturbation-informed signature capable of predicting drug sensitivity in a wide range of untreated tumor cell lines commonly used in cancer research.

Encouraged by these results, we adapted our strategy to mimic a clinical setting utilizing the primary tumor samples of the Cancer Genome Atlas TCGA database (Fig. S7 G). By applying GSVA, we examined the relative distribution of samples from different tumor types based on the calculated sensitivity score, unveiling which cancer types were predicted as generally more susceptible (i.e. kidney renal clear cell carcinoma KIRC, pancreatic adenocarcinoma PAAD, kidney renal papillary cell carcinoma KIRP, glioblastoma multiforme GBM) or less sensitive (i.e. tenosynovial giant cell tumor TCGT, cervical squamous cell carcinoma and endocervical adenocarcinoma CESC, rectum adenocarcinoma READ, colon adenocarcinoma COAD) to **M**, providing a framework for further *in vivo* development of this compound (Fig. 6 E).

Interestingly, the predicted tumor types with the highest and lowest sensitivity turned out to be KIRC and TCGT, respectively, demonstrating that the designed signature was not driven by the original cell type of the cell lines used for its extrapolation and could go beyond the original cancer type. At the same time, PAAD and COAD (pancreatic and colorectal adenocarcinoma, as BxPC-3 and HCT-15 cells) were still among the most and least sensitive, confirming that cell type intrinsic determinants of susceptibility exist and are represented in our signature. Interestingly, intra-tumor variability resulted in a continuous distribution of samples scores within each cancer group, confirming the importance of clinically translating such predictions beyond the tumor type to better address patient-specific therapeutic needs.

To enlarge the accessibility and clinical translatability of our framework, we finally introduced a LASSO regression model to predict the sensitivity of tumor samples in the external reference dataset (Fig. 6 F). We trained a regression model using TCGA basal transcriptomic profiles labelled with the previously predicted sensitivity scores in order to create a self-supervised system able to emulate the prediction irrespective of the context dataset, detaching the predictive tool from the data space. From a clinical perspective, this further step would permit to collect a patient basal transcriptome and feed it to the model, not only improving the performance of the prediction but also avoiding any issue related to data sharing since the model itself does not contain any patients’ sensitive data.

In detail, TCGA samples were labeled according to the calculated continuous sensitivity scores. Next, the model was trained on 80% of the data and tested on the remaining 20%, which efficiently predicted drug sensitivity within the test samples (R = 0.97, RMSE = 0.063) (Fig. 6 G). Notably, such predictive capacity was maintained even when excluding from the transcriptomes all the signature genes used to define the sensitivity score label of the samples, suggesting the biological robustness of the predictive system (R = 0.97, RMSE = 0.066, Fig. S7 H). In fact, while the signature itself was good enough to rank samples based on experimental biological evidence, the model showed to go beyond the initial signature relying on additional predictive features previously not identified.

Overall, we demonstrated how to further employ the integrated RNA-seq and ATAC-seq information to assemble an accurate and clinically-accessible predictive strategy able both to orient drug development and to support the medical practice in the context of precision oncology.

## Discussion

Despite the advances of the last decades, efforts are continuously required to expedite routine use of omic-scale approaches in clinical and pre-clinical settings. Recent work illustrated the potential for omics technologies to accelerate the process of drug discovery from the initial identification of candidate lead compounds up to their pre-clinical and clinical development.^8–14^ Further, improvements in computational approaches for omics data analyses^4, 6, 7^ and an ever-increasing availability of public reference datasets^100^ make it now possible to develop completely new pipelines to address the pharmacological profile of any given drug, from its MoA to sensitivity biomarkers.^1–3^

Here, we combined transcriptome and chromatin accessibility analyses within perturbation experiments to investigate the specific activity profile of 3-CePs, a new class of potential anticancer agents acting as DNA alkylators.^15–21^ Our combined analysis unveiled the basis of the preferential activity of 3-CePs against the pancreatic cancer cell line BxPC-3, which was demonstrated to be unable to properly control proteostasis and DDR under stress conditions upon exposure to the alkylating agents. On the contrary, the low-sensitive colorectal adenocarcinoma cell line HCT-15 potentiated protein folding and catabolism all together activating a more efficient DNA repair after treatment. Due to unresolved genotoxic stress and proteostasis dysregulation, widely described as crosstalking events,^55, 56^ BxPC-3 cells activated the apoptotic branch of the PERK-mediated UPR via CHOP and GADD34, both upregulated after treatment.^61, 62^ Accordingly, such behavior is in line with the described susceptibility of pancreatic cancer adenocarcinoma to ER stress and protein dyshomeostasis.^63^

Beyond validating the described results, the analysis of chromatin accessibility was first employed to identify genes with concordant transcriptional and epigenetic regulation, a step we called *pairwise* integration. Among these genes, we found apoptotic mediators and tumor suppressors upregulated in BxPC-3 and downregulated in HCT-15, as well as redox balance and proteostasis hits upregulated in HCT-15 and downregulated in BxPC-3.

To further evaluate the interaction between transcriptional and chromatin accessibility responses, we proposed here a new versatile approach for the *crosswise* integration of RNA-seq and ATAC-seq, based on vCoCena (vertical Construction of Co-expression network analysis). This approach identified modules of genes co-regulated in the two omic layers across the analyzed experimental conditions. With this standalone method, we not only recapitulated the result of the independent transcriptomic and epigenomic analysis, but we also discovered additional pathways, e.g. actin and TGF β signaling, which modulate the response to the compounds. In detail, actin dynamics were recognized to potentially assist DSBs repair^91^ and a protumorigenic role was established for TGF β in mediating epithelial-mesenchymal transition, both processes that could additionally explain the more efficient response of HCT-15 cells to 3- CePs.^93^ Efficient and versatile, this approach demonstrated to represent a valid option to integrate the information from multi-omic studies substituting the separate examination of each omic dataset.

To further assist the development of 3-CePs, we set up a pilot sensitivity prediction framework readily transferable from the bench to the clinics. We designed a perturbation-informed signature derived from the integrated omic layers filtering the differentially expressed genes between the two cell lines at a steady state for those specifically involved in the cellular response to the treatment. Though based on a limited number of perturbed profiles, this gene signature predicted with high precision the sensitivity to 3-CePs only relying on the untreated transcriptome of test cell lines. The possibility to improve predictions from basal transcriptomes sounds attractive from a clinical perspective since it overcomes the need to screen for thousands of drugs and collect the same amount of profiles from limitedly-available patient samples, such as biopsies.^101^ Applied to TCGA tumor samples, this approach provided a list of susceptible cancer types, e.g. KIRC and PAAD, to support the further development of our drug candidate, and, once transferred on an ML platform, could offer a versatile predictive strategy translatable to the clinics.^6, 102^

In this study, we combined transcriptomic and epigenetic data to guide our exemplary analysis. Nevertheless, the modularity of our framework allows, with only minimal adjustment, its application to other omic technologies or experimental designs. Indeed, the vCoCena integration, which is instrumental for both the biological interpretation of the data and the definition of the perturbation-informed signature, is agnostic of the type of data used as soon as this is reduced to a network of co-regulation.

In conclusion, we present a complete end-to-end workflow to implement the use of multi-omics in drug development, providing a human-readable toolbox to interrogate pharmacological questions in both pre-clinical and clinical settings. We applied this framework to understand the MoA of 3- CePs revealing the cellular determinants of sensitivity to this novel class of drugs and providing precious information for their clinical development as anticancer candidates. Given its versatility, we envision our workflow to be a broadly applicable resource to assist researchers in different steps of the drug discovery and development process.

## Methods

### Cell lines culturing

Colon (HCT-15), pancreatic (BxPC-3), lung (A549) carcinoma cell lines and human embryonic kidney (HEK-293) cells were purchased from ATCC (American Type Culture Collection) while prostate (PC-3) and testis (NTERA-2) carcinoma cell lines were kindly provided by Prof. W. Kolanus (LIMES institute; University of Bonn), neuroblastoma (SH-SY5Y) by Prof. D. Schmucker (LIMES institute; University of Bonn) and epidermoid (A-431) carcinoma by Prof. G. Zunino (Istituto Nazionale dei Tumori di Milano). Cell lines were maintained in logarithmic phase at 37 °C in a 5% carbon dioxide atmosphere using RPMI-1640 (for BxPC-3, HCT-15, PC- 3), DMEM (for A-431, HEK-293, NTERA-2, SH-SY5Y) or Ham’s F-12K (for A549) media (by Gibco or Euroclone) containing 10% fetal calf serum, antibiotics (50 units/mL penicillin and 50 μg/mL streptomycin) and 2 mM L-glutamine (Euroclone).

### Direct detection and quantification of early DNA damage

The extent of early DNA damage induced by 3-CePs in treated cells was assessed by the Fast Micromethod single-strand-break assay. This approach can detect both single and double-strand breaks, as well as alkali-labile adduct sites in the DNA of treated cells. 5,000 cells/well were seeded in 96-well microplates and treated next day for 6 h with **M** (10 nM and 100 nM), **B** (200 nM and 2 µM) or DMSO 0.5%. After treatment, we measured the effect of double and single-strand breaks on the rate of unwinding of cellular DNA in denaturing alkaline conditions by monitoring the fluorescence of a dye that preferentially binds to dsDNA up to 20 min (Pico488 dsDNA quantification reagent, Lumiprobe). The assay was performed following the protocol of Schröder et al.^34^ Two experimental replicates were performed, each one including three technical repeats. Fluorescence signal was acquired by the FLUOstar Omega microplate reader using Omega 5.11 software (BMG LABTECH). The resulting curves based on mean normalized fluorescence values obtained for each treatment and the control (DMSO 0.5%) are reported in Fig. 1 B.

### Cell cycle and flow cytometric H2AX phosphorylation analyses

Possible effects of 3-CePs treatments on the cell cycle distribution of both cell lines were analyzed by FACS, staining cellular DNA with the PI (propidium iodide) dye. In addition, we monitored by antibody staining the phosphorylation of histone H2AX, upstream event of the DDR cascade, after 6 h and 12 h of treatment in order to investigate the ability of BxPC- 3 and HCT-15 cells to detect DSBs. 200,000 cells/well were seeded in 12- well plates and treated next day for 6 h, 12 h or 72 h with **M** (10 nM), **B** (200 nM) or DMSO 0.5%. Cells were harvested, washed with PBS, fixed and permeabilized with the Foxp3 Transcription Factor Staining Buffer Set (eBioscience, cat. #00-5523-00). In detail, cell suspensions were fixed for 1 h at room temperature with FixBuffer, washed twice with PermBuffer and stained with anti-human γH2AX AlexaFluor 488 (Biolegend, clone 2F3, cat. #613405) for 1 h at 4 °C. After the first staining, cells were washed first with PermBuffer, then with PBS and stained secondly with PI (30 min, dark). Samples were acquired on a BD Symphony instrument equipped with 5 lasers (UV, violet, blue, yellow-green, red), the spectral overlap between the channels were determined with single stained samples using FACSDiva (v 9.1.2). Samples were analyzed in FlowJo (BD, v 10.7.1). Events were gated first according to FSC-A and SSC-A and cleaned from cell doubles with 3 consecutive gates (FSC-A vs. FSC-H; SSC-A vs. SSC-H and PI-A vs. PI-H). The frequency of cells within each phase of the cell cycle was calculated using the PI-A signal with the FlowJo built-in algorithm (Watson model with constrained G2 peak). Three biological replicates were obtained per condition and unpaired two-tailed Student’s *t*-test was performed to assess statistical significance (p < 0.05).

### RNA-seq and ATAC-seq experiments

For both RNA-seq and ATAC-seq analyses, 300,000 cells/well were seeded in 6-well plates and treated next day for 6 h, 12 h or 72 h with **M** (10 nM), **B** (200 nM) or DMSO 0.5%. Both for RNA-seq and ATAC-seq samples, three experimental replicates were obtained for each condition.

RNA-seq: at the end of the treatment, cells were washed, resuspended in 1 mL QIAzol reagent (Qiagen) and stored at -80 °C.

ATAC-seq: at the end of the treatment, cells were washed, harvested, resuspended in PBS with EDTA, stained with the LIVE/DEAD Near-IR fixable dye (Invitrogen, cat. #10119) for 10 min at 4 °C, centrifuged and suspended in PBS with EDTA. 20,000 living cells/sample were sorted by FACS and further processed for nuclei isolation and transposition reaction following the protocol of Buenrostro et al.^29^

We extracted the RNA using the miRNeasy mini kit (Qiagen) and checked the RNA integrity and quantity using the tapestation RNA assay on a tapestation4200 instrument (Agilent). We used 750ng total RNA to generate NGS libraries using the TruSeq stranded total RNA kit (Illumina) following manufacturer’s instructions and generated ATAC-libraries from tagmented cells following the protocol of Buenrostro et al. In both cases we checked library size distribution via tapestation using D1000 (RNA) and D5000 assays (ATAC) respectively on a Tapestation4200 instrument (Agilent) and quantified the libraries via Qubit HS dsDNA assay (Invitrogen). We clustered the libraries at 250pM final clustering concentration on a NovaSeq6000 instrument using SP and S2 v1 chemistry (Illumina) and sequenced paired-end 2*50 cycles before demultiplexing using bcl2fastq2 v2.20.

### RNA-seq data analysis

Reads were aligned and quantified with STAR (v 2.5.2a)^103^ using standard parameters and mapped against the GRCh38p13 human reference genome (Genome Reference Consortium). Raw counts were imported, pre-filtered to exclude low-count genes (<100 reads, 17.693 mapped transcripts), normalized and VST-transformed (variance stabilizing transformation) following the DESeq2 (Bioconductor, v 1.26.0) pipeline using default parameters.^104, 105^ SVA (surrogate variable analysis) was applied to identify latent variables responsible for batch effects and four of them were included in the DESeq2 model.^106^ All present transcripts were used as input for principal component analysis (PCA). The call for differentially expressed genes was performed for all treated vs control comparisons (separate cell lines) using an adjusted p-value threshold equal to 0.05, where IHW (IHW: independent hypothesis weighting) was adopted for multiple testing. Only protein-coding hits were considered for further functional analyses on DE genes. GSEA (gene set enrichment analysis) based on the GO (gene ontology) *biological process* database was employed for functional enrichments, both based on DE genes (Supplementary data 1, 2) or log2FC-based ranks. All enrichment dotplots report the Count and p-value associated with each term, when p < 0.05.

Representative enrichment terms in Fig. 2 C were selected manually from enrichment maps obtained for each group of genes depicted in the dotplot (Supplementary data 7): to remove semantic redundancy, only the most significant nodes among those converging into the same hub were reported (higher Count and lower p-value, example in Fig. S2 F). SVA batch-corrected normalized vst-transformed counts were used as input for boxplots, heatmaps and log2FC-based GSEA. Hierarchical clustering was applied to identify blocks of DE genes with similar regulations across conditions as reported in the presented heatmaps (Fig. 2 D, S3 A). In the same heatmaps, row-scaled expression levels of cell-specific DE genes elicited at 6 h and 12 h were reported separately for each of the analyzed conditions.

### ATAC-seq data analysis

After adapter trimming using Trimmomatic v 0.36^107^, the sequencing reads were aligned bowtie2 v 2.3.5 against the GRCh38p13 human reference genome.^108^ Subsequently, duplicated reads were removed using Picard *dedup* function and the transposase-induced offset was corrected using the deeptools v 3.1.3 *alignmentSieve* function.^109^ After sorting and indexing bam files with samtools v 1.9.,^110^ peak calling was performed using MACS2 v 2.1.2.^111^ Peak regions from sample-specific peak calling results were unified in R v 3.6.2 using the *reduce* function implemented in the GenomicRanges package v 1.38.0.^112^ prior to quantification of sequencing reads in these unified peak regions using the *summarizeOverlaps* function implemented in the GenomicAlignments package v1.22.1.^112^ Raw counts were pre-filtered to exclude low-count peaks (<20 reads, 63.434 mapped peaks), normalized and VST-transformed following the DESeq2 (Bioconductor, v 1.26.0) pipeline using default parameters.^104, 105^ Peak regions were annotated using *ChIPseeker* v1.22.1. All present peaks were used as input for principal component analysis (PCA). The call for differentially accessible regions (DARs) was performed for all treated vs control comparisons (separate cell lines) considering a p < 0.05 threshold. Only peaks mapping in promoters of protein-coding regions were considered for further functional analyses. GSEA (gene set enrichment analysis) based on the GO (gene ontology) *biological process* database was employed for functional enrichments based on DAR-associated genes (Supplementary data 3). Normalized and vst-transformed counts were used as input for heatmaps and boxplots. Hierarchical clustering was applied to identify blocks of DAR-associated genes with similar regulations across conditions as reported in the presented heatmaps (Fig. 4 C, S5 D). In the same heatmaps, row-scaled accessibility levels of cell-specific DARs at 6 h and 12 h were reported separately for each of the analyzed conditions. For the *pairwise* integration between transcriptional and chromatin accessibility data, we identified hits having the same sign of regulation in RNA-seq and ATAC-seq which were DE (protein-coding) and/or DAR-associated (protein-coding mapping in promoters). Since a delay could exist between a prior chromatin remodeling and a detectable variation in the respective transcript level, pairwise comparisons were considered not only at the same time point in both omic layers but also between chromatin changes at 6 h and transcriptional responses at 12 h. We reported in Fig. 4 D only hits with (log2FCRNA-seq + log2FCATAC-seq) > 1 or < -1. Interesting gene names for each of the considered comparisons were also reported.

### Crosswise integration of RNA-seq and ATAC-seq data

The *crosswise* integration of transcriptomic and chromatin accessibility data was achieved through an adaptation of the CoCena (construction of co-expression network analysis - automated) tool, which can identify modules of genes showing similar regulation across conditions of interest. The core principles driving both network construction and gene modules detection by CoCena have been described previously.^88^ In this analysis, we first optimized the design of separate co-expression networks for the RNA-seq and ATAC-seq layers. To avoid the creation of networks mostly describing cell type differences, we calculated separate networks for BxPC-3 and HCT-15 cells which were then integrated horizontally through hCoCena^2^.^88^ The union of all DE and promoter DAR-associated genes detected in treated conditions was selected as input for constructing all networks. For the construction of cell-specific networks, the specified Pearson correlation cutoffs, edges and nodes for RNA-seq (BxPC-3: cutoff=0.801, edges=356851, nodes=4266; HCT-15: cutoff=0.772, edges=154497, nodes=4321) and ATAC-seq (BxPC-3: cutoff=0.702, edges=48280, nodes=3479; HCT-15: cutoff=0.733, edges=13336, nodes=3350) were used. The horizontally integrated networks contained the union of all nodes and edges coming from parent networks, where edges between nodes connected in both parent layers were recalculated as a mean of their original weights. Clustering of the resulting RNA-seq and ATAC-seq networks was performed based on the *infomap* algorithm, where a threshold of minimum of 15 nodes per cluster was applied (Fig. S6 B, C).^113^

Subsequently, inter-omic integration by vCoCena was applied to construct the final network. In this case, the correlation between the mean group-fold change (GFC) pattern of modules belonging to the two layers was calculated to identify clusters of genes with similar regulation, suitable for *crosswise* integration. Edges from the two separate networks were selected for contributing to the integrated one based on a minimum cross-layer correlation which could guarantee the maximum mixture between layers in identified module pairs (minimum correlation cutoff=0.73, edges=628783, nodes=8067). The new network was reclustered exploiting again the *infomap* algorithm, applying a higher threshold of a minimum of 30 nodes per cluster, and mean GFCs were recalculated: the resulting integrated modules included nodes originally derived from the two separate layers in different ratios, as shown in the relative heatmap (Fig. 5 B). GO-based GSEA was performed on detected modules of genes (Supplementary data 4) and the most significant terms (p < 0.05) were reported.

### Sensitivity signature construction and prediction pipeline

For the signature of sensitivity to **M**, relevant modules from the crosswise vCoCena integration were selected as follows (Fig. S7 A): for each module, in both cell lines separately, we calculated the difference between the GFC (group fold-change) of the control and the **M** 6 h treated groups (ΔGFC(cell line) = GFC(**M**6h) - GFC(ctrl)). The early time point was selected to guide the signature construction since from upstream analyses it turned out to be the most informative of cell responses to 3-CePs. The threshold score was then calculated as the difference between the previously obtained ΔGFCs for the two cell lines (thrscore = ΔGFC(BxPC-3) - ΔGFC(HCT-15)). Modules with thrscore above q50, thus modules where the regulation was sufficiently different in the two cell lines after treatment with **M**, were selected (*cyan*, *darkgreen*, *darkgrey*, *darkorange*, *gold*, *indianred*, *khaki*, *lightgreen*, *steelblue*, *orchid*). Genes from the identified modules were grouped together and further considered to drive the definition of our signature of interest.

Further on, DE analysis was performed between BxPC-3 and HCT-15 untreated control groups to identify baseline DE genes up- and downregulated in the high-sensitive cell line (log2FC threshold equal to 1, padj < 0.01). In fact, given the much higher availability and clinical spendability of RNA-seq compared to ATAC-seq profiles, the signature was finally constructed only from basal transcriptomes. In particular, we further selected among the identified module genes only those that were also DE between the two untreated controls, ending up with a restricted group of genes showing compound- and cell line-specific regulation upon perturbation but, meanwhile, a significantly different basal expression in BxPC-3 cells. This perturbation-informed signature was composed by 294 genes upregulated (signature *up*) and 170 genes downregulated (signature *down*) in the high-sensitive BxPC-3 cells (listed in Supplementary data 5).

To validate the predictive performance of the obtained signature, GSVA (gene set variation analysis) was performed both with *up* and *down* signatures on the basal RNA-seq profiles of cancer cell lines included in the HPA (Human Protein Atlas).^99^ A sensitivity score was calculated for each cell line as the difference between the ESs (enrichment scores) of the *up* and the *down* signatures. The predicted rank was validated on selected cell lines (A-431, A549, HEK-293, NTERA-2, PC-3, SH-SY5Y) as described in the next paragraph and Pearson correlation between predicted sensitivity scores and viability decrease in cells treated with **M** 10 nM for 72 h was calculated. Two control signatures of the same size were also tested: 1) a random genes signature (composed by random genes among those annotated in the RNA-seq profile of HPA cell lines) 2) a control signature composed by the top up- and down- log2FC DE genes between the two cell lines (listed in Supplementary data 6).

GSVA (v 1.38.2) was applied also on basal transcriptomes of samples from the Cancer Genome Atlas TCGA database and their sensitivity score was calculated as previously indicated. The relative distribution of samples from different tumor types in terms of calculated sensitivity score was plotted in Fig. 6 E, together with the indicated median value for each group, to identify possibly more susceptible tumor types.

Finally, the signature-based prediction was used to train a LASSO-based classifier (*cv.glmnet* function in *glmnet* package v 4.1 to assess lambda penalty, *predict* function in *stats* package v 4.0.3 for actual prediction). Briefly, TCGA samples were assigned to a continuous label based on the previously inferred sensitivity scores. We next trained the classifier with 80% of these profiles and tested it on the remaining 20%: Pearson correlation and RMSE were calculated to evaluate the predictive performance of the classifier. To assess the biological robustness of our signature and of the obtained model, the classifier was trained and tested also using transcriptomes cleaned up from genes belonging to our signature.

### Validation of 3-CePs sensitivity prediction on cancer cell lines

The rank of sensitivity to **M** obtained from the newly constructed signature was validated on a subset of available cell lines included in the Human Cell Atlas. The selected cell lines spanned quite well between the max and min detected susceptibility scores. Here are the screened cell lines from the one predicted as most sensitive: PC-3, A549, A-431, SH-SY5Y, NTERA-2, HEK-293. 5,000 cells/well were seeded in 96-well microplates and after 24 h treated with **M** 10 nM for 72 h. Cell viability was assessed at the end of the treatment by MTT, following previously adopted protocols.^19^ Mean values of residual viability and standard deviations obtained from two independent experiments in duplicated microplates, each one containing three technical replicates, are reported in Table S1. Pearson correlation between mean residual viability and predicted susceptibility score in considered cell lines was calculated and reported in Fig. 6.

### Statistics and reproducibility

Sample size was defined empirically to ensure robust statistical analysis. Unpaired two-tailed Student’s *t*-test was performed to assess statistically significant differences (p < 0.05) in cell cycle and H2AX phosphorylation analyses between treated and control conditions (*n*=3). All correlation coefficients were calculated with a Pearson’s test. The adopted statistical tests, the considered significance levels and the number of biological replicates are also reported in figure legends. Box plots are in the style of Tukey, where the center of the box represents the median of values, hinges represent the 25th and 75th percentile and the whiskers are extended not further than the 1.5 * IQR (inter quartile range). The analysis was performed on R (v. 3.6.2 or 4.0.3): the specific packages used for the analysis, their version and relevant parameters used are explained in the *Methods* sections. All plots were generated with *ggplot* (v. 3.3.2) except for the heatmaps which were generated with the R package *complexheatmap* (v. 2.2.0). To ensure the reproducibility of the manuscript results, all the analyses were conducted within a containerized environment (Docker). RNA-seq and ATAC-seq analyses were performed with the docker image jsschrepping/r_docker:jss_R362 (https://hub.docker.com/r/jsschrepping/r_docker). The rest of the analysis was conducted with the image lorenzobonaguro/cocena:v3 (https://hub.docker.com/r/lorenzobonaguro/cocena) for compatibility with the CoCena pipeline.

### Data Availability

All raw data included in this study are available at gene expression omnibus (GEO). Raw RNA-seq data and count matrix under the GEO accession number GSE179057. Raw ATAC-seq data and peak matrix are available under the accession number GSE179059. Both datasets are collected in a GEO SuperSeries (GSE179064).

### During the review process reviewer can access the private dataset at the link: https://www.ncbi.nlm.nih.gov/geo/query/acc.cgi?acc=GSE179064 using the provided access token

The cell line expression data employed in the prediction pipeline were downloaded from https://www.proteinatlas.org/about/download. The file *RNA HPA cell line gene data* contains transcript expression levels summarized per gene in 69 cell lines and is based on the Human Protein Atlas version 20.0 and Ensembl version 92.38.

Similarly, the TCGA expression data from cancer cell samples (the Cancer Genome Atlas) were downloaded from the same web page of the Human Cell Atlas (*Transcript expression levels summarized per gene in 7932 samples from 17 different cancer types*). Data are based on The Human Protein Atlas version 20.0 and Ensembl version 92.38.

## Code availability

The code to reproduce both pre-processing and downstream analyses reported in this manuscript will be made publicly available on GitHub upon acceptance. The CoCena script is accessible at https://github.com/Ulas-lab/CoCena2.

## Author contributions

Conceptualization was by C.C., L.B, S.M., V.G., R.G., A.C.A., J.L.S and B.G. The methodology was devised by C.C., L.B., J.S.-S., M.O, S.W.-H., T.U., while C.C, L.B., A.H., T.H., M.D.F., K.H. performed formal experiments and C.C., L.B., J.S.-S. the data analyses. C.C, L.B. carried out the investigations. The manuscript was written by C.C, L.B., J.L.S and B.G. while all authors reviewed and edited it. The project was supervised by C.C., L.B, J.L.S and B.G. Resources were provided by V.G., J.L.S., B.G.

## Correspondance

Correspondence to J. L. Schultze and B. Gatto.

## Supplementary Information

**Figure S1.**
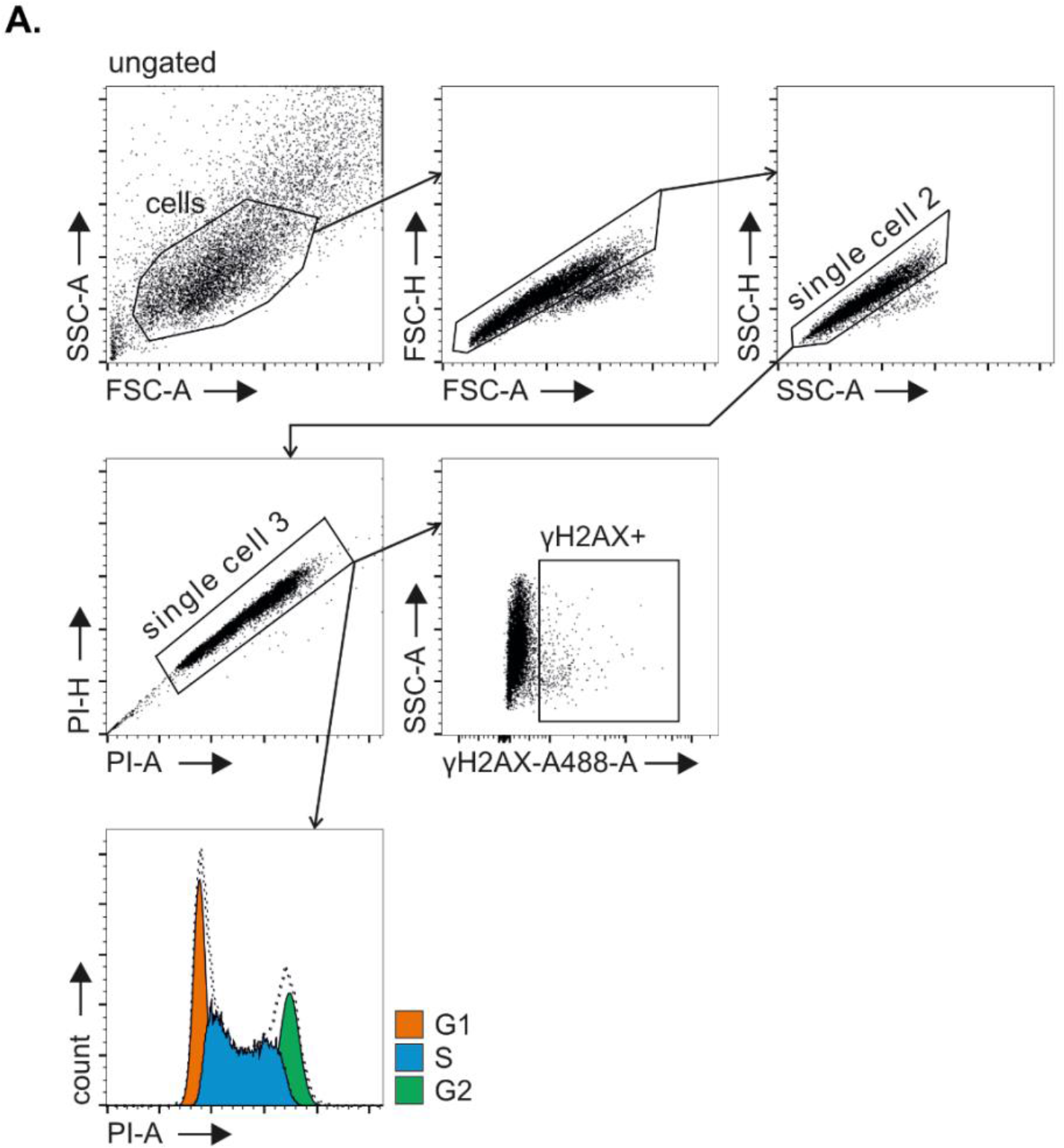
**A** Flow cytometry gating strategy for the cell cycle analysis and γH2AX induction reported in Fig.1 C and 1 D.

**Figure S2.**
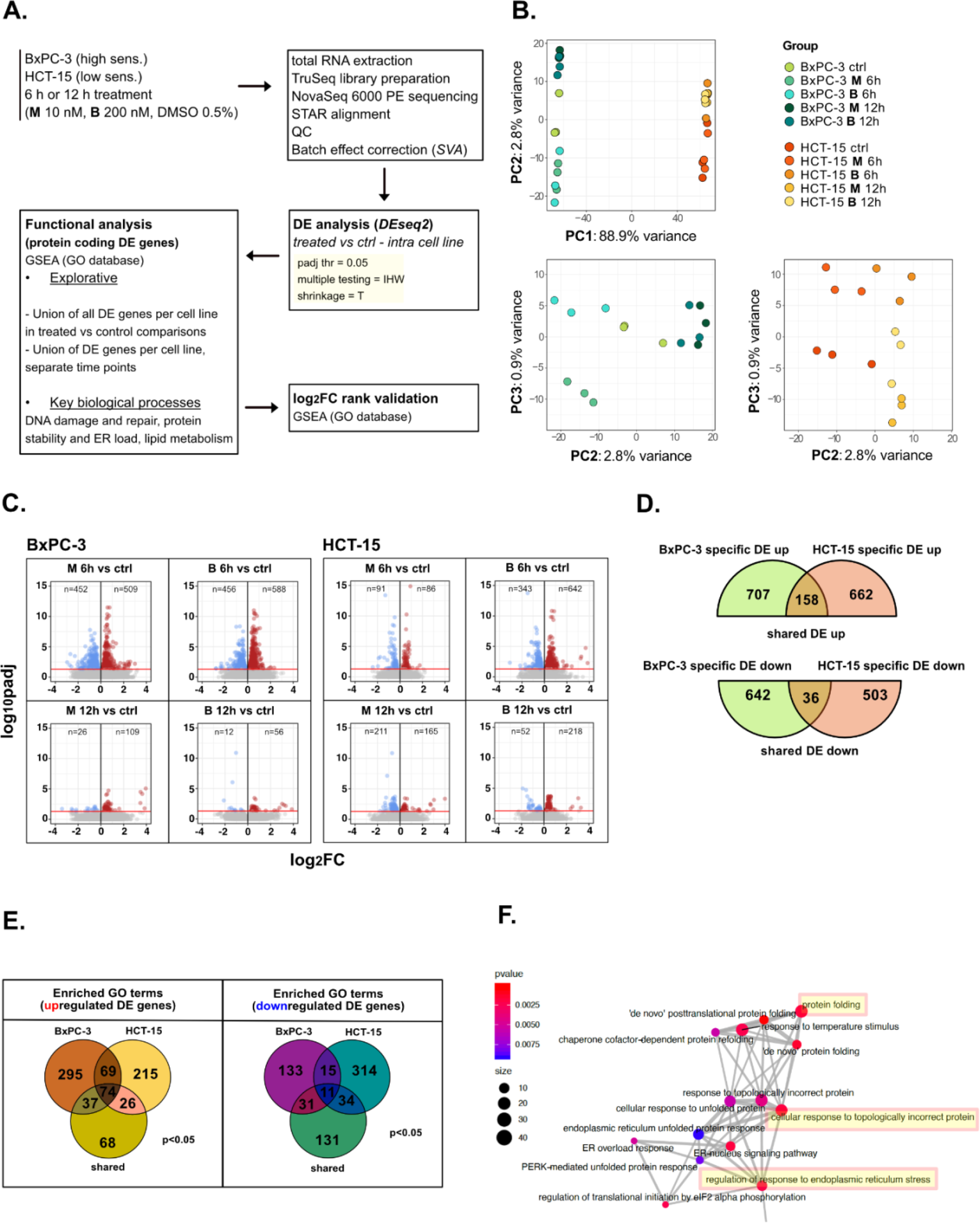
**A** Scheme of the applied workflow for the RNA-seq analyses. **B** Principal component analysis (PCA) post SVA batch correction of RNA-seq data: PC1 vs PC2 showed sample separation by cell line, PC2 vs PC3 (cell lines depicted separately) showed treatment and time point separation. **C** Volcano plots reporting up- and downregulated DE genes in all treated vs control comparisons (adjusted p-value < 0.05). **D** Venn plot reporting the number of specific and shared up- and downregulated DE genes between BxPC-3 and HCT-15 cells (union of DE genes in all treated vs control comparisons). **E** Enriched GO terms (p < 0.05) derived from GSEA on BxPC- 3 and HCT-15 DE genes (union of DE genes in all treated vs control comparisons) and on shared DE genes (up- and downregulated separately). **F** Schematic representation of enrichment map-based selection of representative GO terms to be reported in Fig. 2 C.

**Figure S3.**
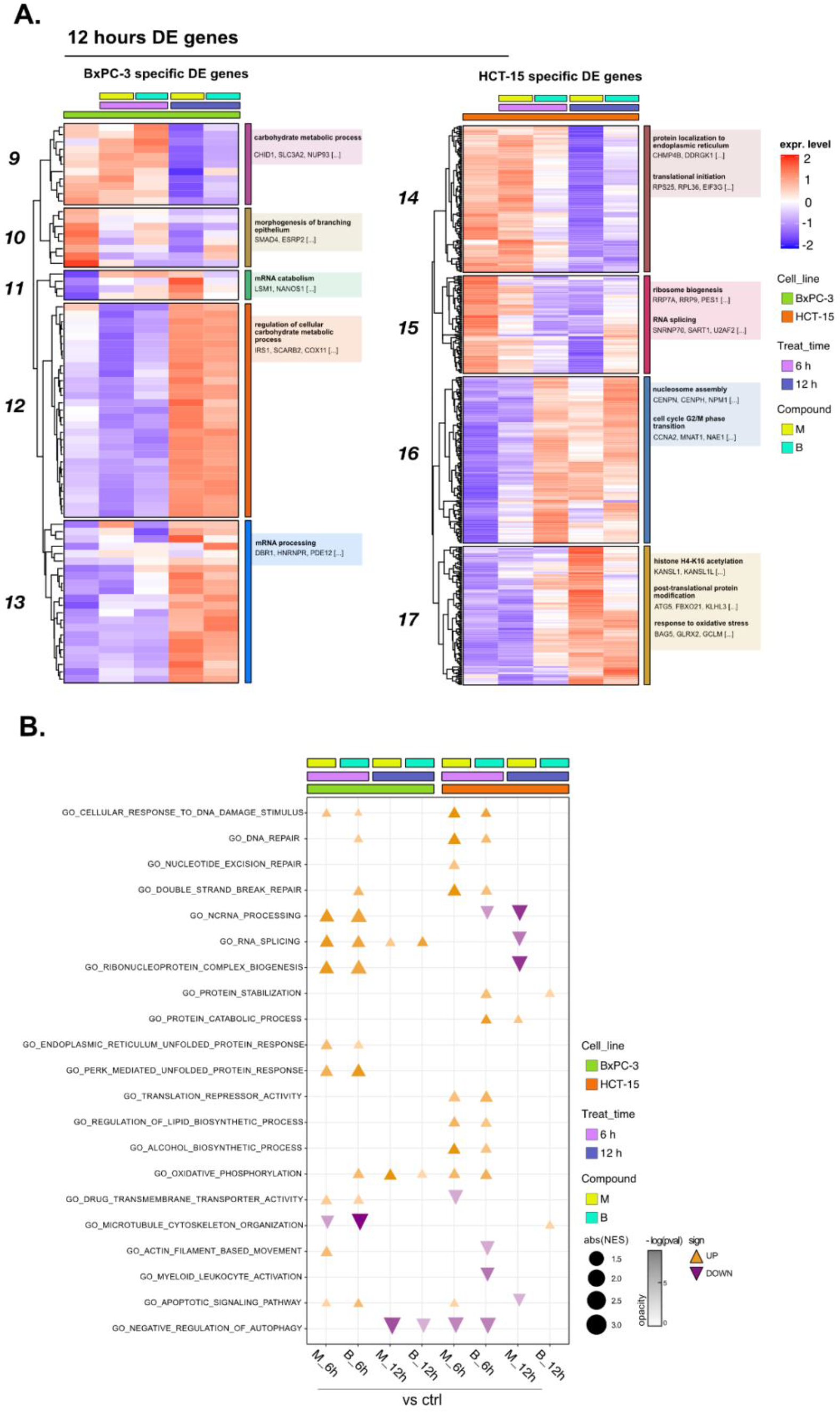
**A** Expression level of cell-specific 12 h DE genes across test conditions. GSEA was performed on modules with similar regulation identified by hierarchical clustering: for each cluster, representative GO terms and genes of the associated load are reported. **B** GO database functional enrichment (GSEA) obtained from log2FC ranks in all treated vs control comparison both in BxPC-3 and HCT-15 cells. For each identified biological process, enrichments in terms of absolute normalized enrichment score (abs(NES)) and -log(p) of representative terms are reported (p < 0.05).

**Figure S4.**
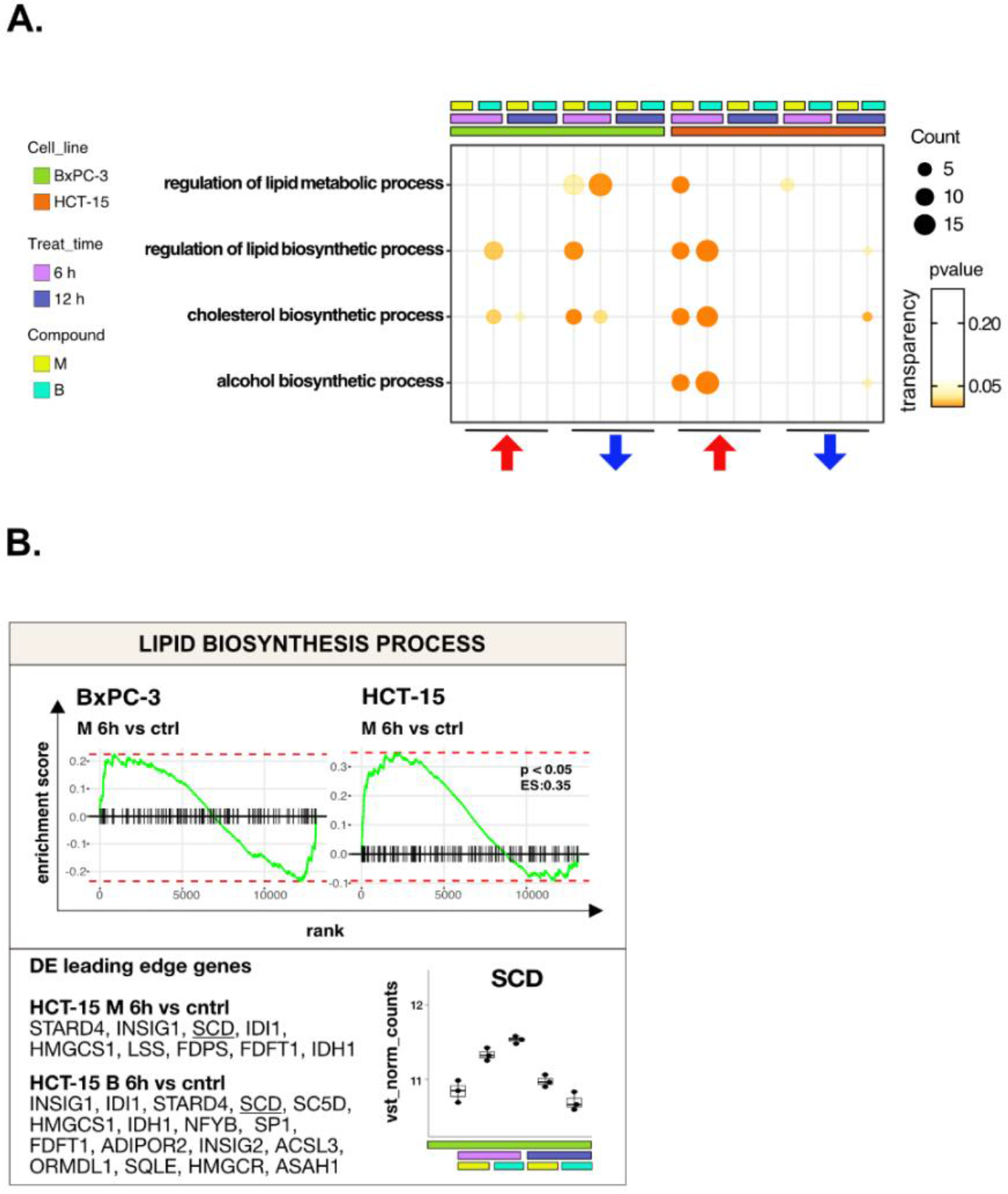
**A** GSEA for terms related to *lipid metabolism* performed on DE genes detected in each of the considered treated vs control comparisons. For each GO term (p < 0.05), enrichments in terms of Count and p-value are reported. **B** GSEA enrichment plots for the *lipid biosynthesis process* pathway obtained from log2FC ranks for each of the considered treated vs control comparisons. DE leading edge genes are also reported, together with boxplots showing the expression level of SCD (vst-transformed normalized counts) in BxPC-3 cells.

**Figure S5.**
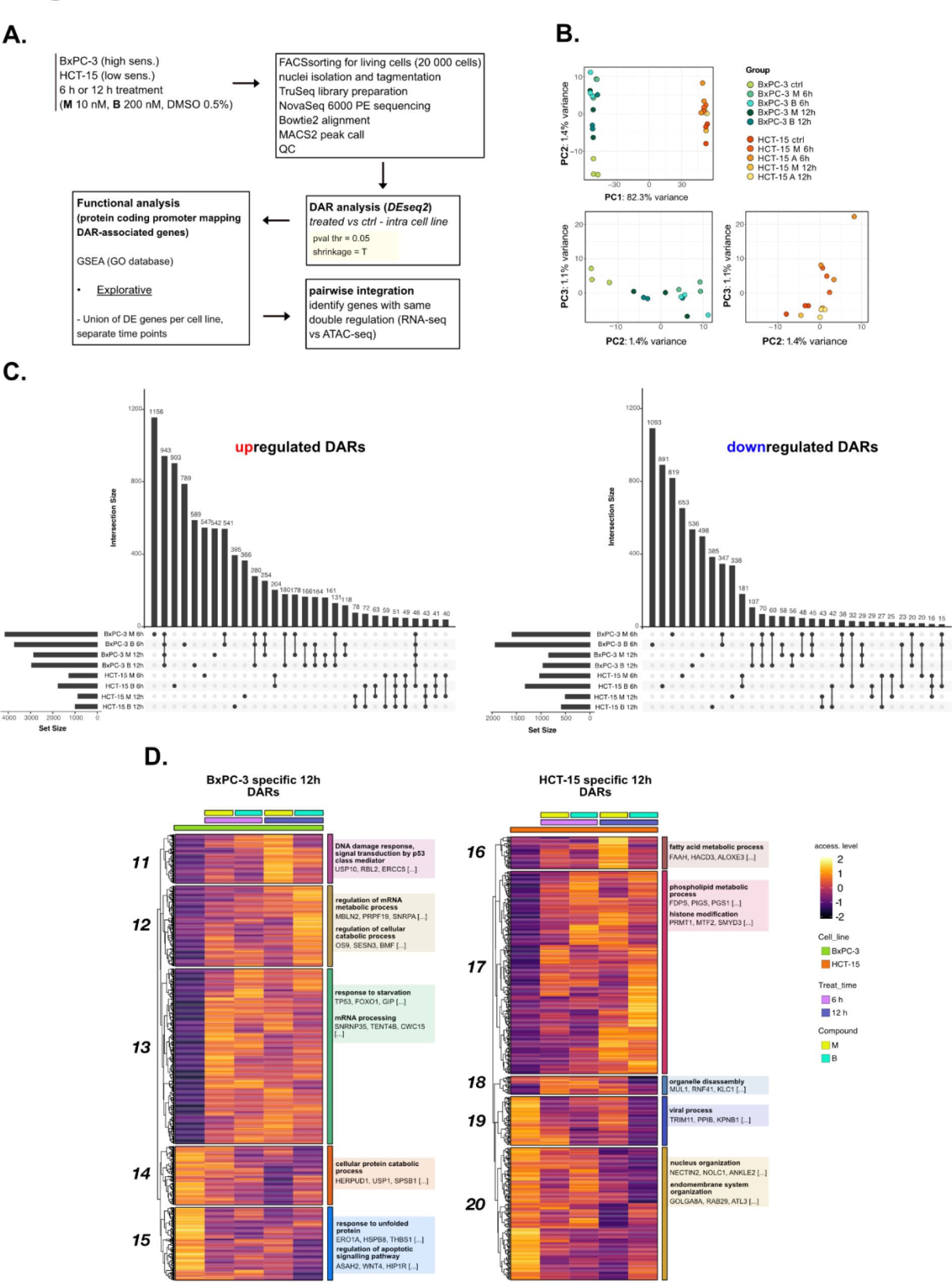
**A** Scheme of the applied workflow for the ATAC-seq analyses. **B** Principal component analysis (PCA) of ATAC-seq data: PC1 vs PC2 showed samples separation by cell line, PC2 vs PC3 (cell lines depicted separately) showed treatment and time point separation. **C** Upset plots reporting up- and downregulated DARs (p < 0.05) and their overlap between all treated vs control comparisons in both cell lines. **D** Accessibility level of cell-specific 12 h DARs across test conditions. GSEA was performed on genes associated with DARs with similar regulation, grouped in modules identified by hierarchical clustering: for each cluster, representative GO terms and genes of the associated load are reported.

**Figure S6.**
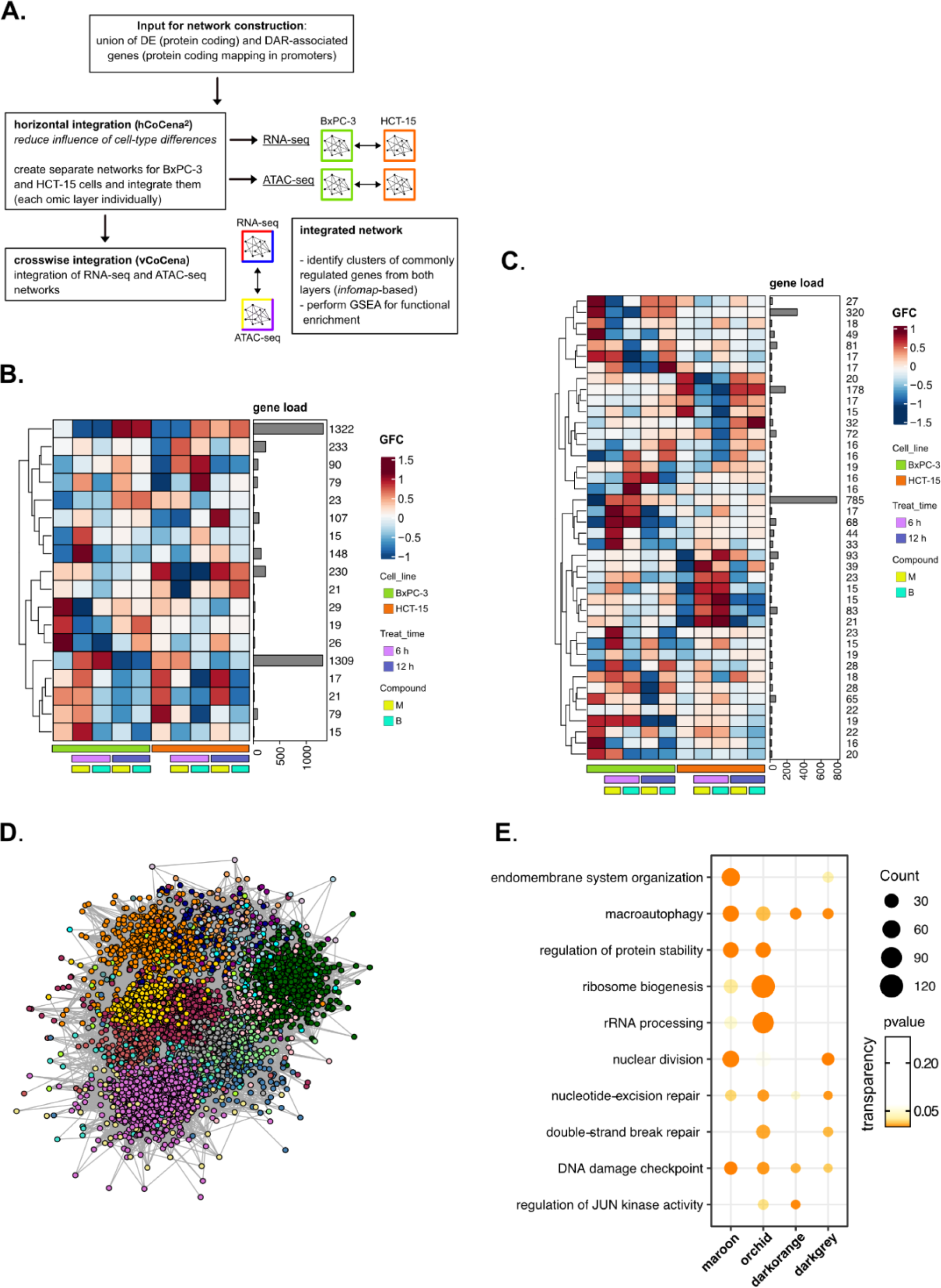
**A** Scheme of the applied workflow for the *crosswise* integration analysis. **B** Horizontally integrated modules of genes from the RNA-seq layer and associated GFC (group fold change) pattern of regulation across conditions. **C** Horizontally integrated modules of genes from the ATAC-seq layer and associated GFC (group fold change) pattern of regulation across conditions. **D** *Crosswise* integrated vCoCena network. **E** Most representative GO terms from GSEA on genes of the *maroon, orchid, darkorange, darkgrey* modules. For each GO term (p < 0.05), enrichments in terms of Count and p-value are reported.

**Figure S7.**
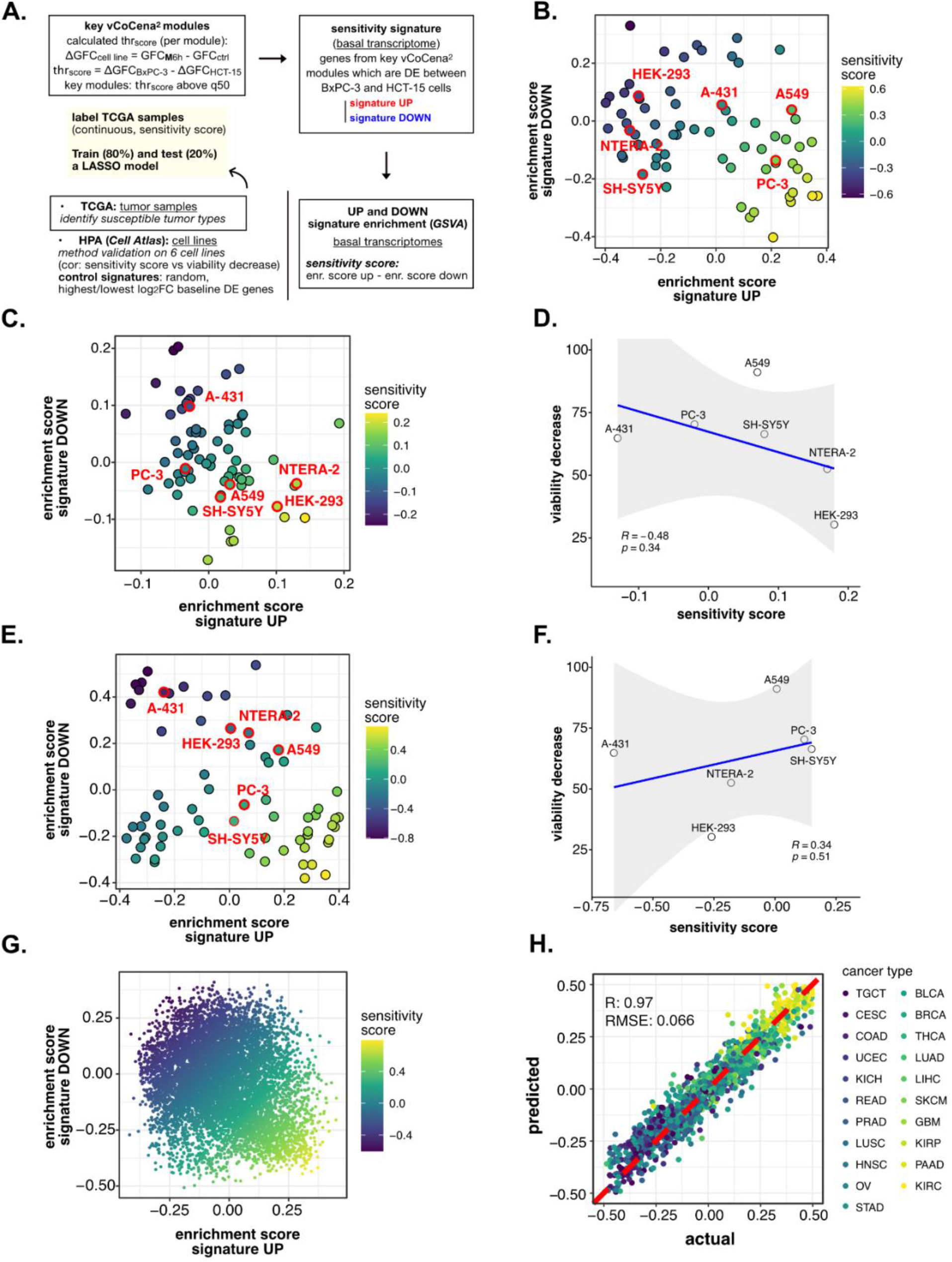
**A** Scheme of the applied workflow for the sensitivity signature construction and associated drug susceptibility prediction. **B** HPA (*Cell Atlas*) cell lines separation based on GSVA enrichment of our newly constructed up vs down signatures of sensitivity to M. Color scale reflects samples predicted sensitivity score (up signature enrichment - down signature enrichment). **C** HPA (*Cell Atlas*) cell lines separation based on GSVA enrichment of random up vs down signatures. Color scale reflects samples predicted sensitivity score (up signature enrichment - down signature enrichment). **D** Pearson correlation between predicted sensitivity score and viability decrease in a subset of HPA (*Cell Atlas*) cell lines (validation set) using a random signature. **E** HPA (*Cell Atlas*) cell lines separation based on GSVA enrichment of control up vs down signatures. GSVA was performed using a control signature composed by DE genes with top up and down log2FC. Color scale reflects samples predicted sensitivity score (up signature enrichment - down signature enrichment). **F** Pearson correlation between predicted sensitivity score and viability decrease in a subset of HPA (*Cell Atlas*) cell lines (validation set) using a control signature composed by DE genes with top up and down log2FC. **G** TCGA tumor samples separation based on GSVA enrichment of our newly constructed up vs down signatures of sensitivity to M. Color scale reflects samples predicted sensitivity score (up signature enrichment - down signature enrichment). **H** Predictive performance after the exclusion of genes belonging to our signature from training and test set transcriptomes (Pearson correlation R and RMSE are reported).

**Table S1.** Average viability decrease in cell lines treated with **M** 10 nM for 72 h with associated standard deviation (SD). For each cell line, predicted sensitivity scores based on our perturbation-informed signature (signature SS), a random one (random SS), a control one based on top up and down log2FC DE genes between BxPC-3 and HCT-15 (topFC SS) were also reported.

| Cell line | Viability decrease | SD | signature SS | random SS | topFC SS |
| --- | --- | --- | --- | --- | --- |
| HEK-293 | 30,3 | 15,1 | -0,37 | 0,18 | -0,26 |
| NTERA-2 | 52,5 | 6,14 | -0,28 | 0,17 | -0,18 |
| SH-SY5Y | 66,4 | 4,61 | -0,08 | 0,08 | 0,15 |
| A-431 | 64,8 | 7,54 | -0,03 | -0,13 | -0,66 |
| A549 | 91,1 | 2,85 | 0,23 | 0,07 | 0,007 |
| PC-3 | 70,3 | 0,14 | 0,35 | -0,02 | 0,12 |

SD: standard deviation
signature SS: perturbation-informed signature sensitivity score
random SS: random signature sensitivity score
topFC SS: top up and down log<sub>2</sub>FC control signature sensitivity score

